# *SenePy*: Unveiling the Cell-Type Specific Landscape of Cellular Senescence through Single-Cell Analysis in Living Organisms

**DOI:** 10.1101/2023.08.30.555644

**Authors:** Mark A. Sanborn, Xinge Wang, Shang Gao, Yang Dai, Jalees Rehman

**Author notes:** Please address correspondence to: Jalees Rehman, MD, Department of Biochemistry and Molecular Genetics, The University of Illinois, College of Medicine, 900 S. Ashland Ave (M/C 669), Chicago, IL, 60607.

## Abstract

Senescent cells accumulate in tissues with organismal age and contribute causally to multiple chronic diseases. *In vivo* senescent cell phenotypes are heterogeneous because cellular context and stressors vary by cell type and tissue. Due to the variability of senescence programs, there is no universal method to identify senescent cells and even widely used markers, such as *CDKN2A*, are not ubiquitous. Therefore, we interrogated the *Tabula Muris Senis* mouse single-cell aging atlas and an array of single-cell datasets from human donors that spanned many ages to find cell-specific signatures of cellular senescence. We derived 75 mouse and 65 human senescence signatures from individual cell populations. *CDKN2A* and other markers of senescence were overrepresented in these signatures but there were many novel senescence genes present at higher rates. Within individual cell populations, we observed multiple programs of senescence with distinct temporal and transcriptional characteristics. We packaged the signatures along with a single-cell scoring method into an open-source package: *SenePy. SenePy* signatures better recapitulate cellular senescence than available methods when tested on multiple *in vivo* RNA-seq datasets and a p16^ink4a^ reporter single-cell dataset. We used *SenePy* to map the kinetics of senescent cell accumulation across 97 cell types from humans and mice. *SenePy* also generalizes to disease-associate senescence and we used it to identify an increased burden of senescent cells in COVID-19 and myocardial infarction. This work provides a significant advancement towards our ability to identify and characterize *in vivo* cellular senescence.

## INTRODUCTION

Aging is a key risk factor for many chronic diseases^1^. One biological manifestation of organismal aging is cellular senescence (CS), a phenomenon characterized by permanent cell cycle arrest, impaired homeostatic cellular function, and activation of the senescence-associated secretory phenotype (SASP) which involves the release of pro-inflammatory proteins, proteases, and other bioactive paracrine factors^2^. Senescent cells accumulate in tissues with increasing organismal age, but senescent cells are found even in young organisms and can accrue prematurely due to exogenous stressors^3, 4^. Accumulated senescent cells contribute to sterile inflammation, tissue remodeling, and local dysfunction which ultimately drives various pathologies^5^. CS contributes to a wide array of chronic diseases, including cardiovascular disease, neurodegeneration and diabetes^6-8^. The senescent cell burden in aged organisms also contributes to runaway inflammation and poor outcomes in acute diseases, such as those from coronavirus infection^9^. Targeted clearance of senescent cells with senolytics can mitigate disease severity and increase healthspan^7-10^, but in some contexts their elimination may exacerbate disease^11^. Despite the growing understanding of the role of cellular senescence in aging and various diseases, *in vivo* cellular senescence remains poorly phenotypically and mechanistically characterized^2, 12^. The majority of CS markers have been identified in cultured cells subjected to experimental conditions that may not accurately represent a living system. More comprehensive sets of markers are required to robustly study cellular senescence in living systems.

One of the biggest challenges in studying *in vivo* cellular senescence is the high degree of heterogeneity in which it presents^2^. Senescence has been observed in numerous cell types across all major organs. Senescent cells partially lose their pre-senescence identities and phenotypes but this suggests that the mechanistic paths to the senescent states varies between cell types^13, 14^. Cells are also subject to various cell-intrinsic and extrinsic stressors which drive senescence. Telomere attrition is a well-known senescence trigger, but telomere-independent DNA damage, oxidative stress, and oncogenic signaling can also induce cellular senescence^5^. Paracrine factors released in the SASP state and cell-surface signaling from senescent cells can induce secondary senescence among otherwise healthy cells in close proximity^15, 16^. Drivers of senescence shift the transcriptional landscape of senescent cells^13, 14^, but this has been primarily studied in cultured cells. The tissue context or cell identity-specific transcriptional landscapes of cellular senescence have not been fully defined. For example, it is not clear whether the senescence transcriptional programs in skin fibroblasts exposed to UV light differ from the senescence programs of fibroblasts in internal organs protected from light. The heterogeneity arising from different stressors, tissues, and cell types makes it difficult to broadly apply transcriptional signatures of senescence derived from *in vitro* cultured cells which have been removed from their *in situ* tissue environment. There is no universal signature or marker of CS exists^2^. Even the cell cycle arrest inducer p16^ink4a^, which is widely accepted as one of the most specific markers of cellular senescence, is not always required for senescence induction and its use as a sole marker in transcriptomics data is confounded by the fact that the corresponding *CDKN2A* locus encodes for multiple genes with overlapping sequence identity^17-19^. Other markers of senescence may be constitutively expressed in some cell types or upregulated in general with organismal age or inflammation. Recent work has utilized literature screening and transcriptomics to find a gene set that is broadly differentially abundant in senescent cells^20^, but this does not account for tissue-, cell-, or stress-specific heterogeneity and may not capture all programs of cellular senescence. There remains a need to identify and characterize tissue- and cell-specific cellular senescence programs^21^.

In this study, we aggregate and interrogate large-scale single-cell RNA-sequencing datasets across tissues and ages, both in mice and humans to define *in vivo* cellular senescence heterogeneity. We developed a novel algorithmic approach to identify cell-type-specific senescence signatures. We used a p16^ink4a^ reporter mouse model dataset along with other transcriptomics datasets to validate our approach. We have generated the open-source Python package *SenePy* (https://github.com/jaleesr/senepy). *SenePy* allowed us to map the kinetics of cellular senescence in many tissues and cell types with respect to organismal age and in the context of disease. Using *SenePy* we were able to identify senescent cells across several tissues and examine similarities as well as tissue-specific and cell-type-specific signatures of cellular senescence.

## RESULTS

### Known senescence markers are cell-type-specific and poorly characterize *in vivo* cellular senescence

We examined the expression of established CS markers in comprehensive mouse and human single-cell atlases to determine their dynamics with age and their cell-type-specificity. For mouse data, we utilized the *Tabula Muris Senis*^*22*^ resource which is comprised of 328K cells from 19 tissues collected between 1 and 30 months of age (**Sup Fig 1A**). The human data utilized in this study are from 7 studies^23-29^ that span 37 tissues from individuals ranging in age from 1 to 92 years old, altogether comprising 1.6M cells (**Sup Fig 1B**). We took the union of *SenMayo*^20^ (n = 125), which is a recently published list of senescence-associated genes, and a novel curated senescence gene set (n = 110) to establish a panel of 181 experimentally validated CS maker genes (∩= 52, hypergeometric p = 6.4x10^-94^). This panel of known or validated senescence markers served as a starting point for our downstream analyses.

To assess how this set of senescence markers overlapped with cell-specific and universal organismal aging genes in mice, we compared them to a study that identified 76 cell-specific signatures and a universal aging signature (n = 330 genes) from *Tabula Muris Senis*^*30*^ (**Fig 1A**). Only the CS markers *Cd9, Ctnnb1, and Jun* were present in the universal organismal aging signature (defined by genes upregulated with age in half of the cell types), which can be explained by random chance (hypergeometric p = 0.58). *CDKN2A* (p16^ink4a^ encoding gene), which is widely accepted as one of the most universal and specific markers of CS, was not present in any of the cell-specific organismal aging signatures. Furthermore, only 15 of the 76 cell-specific aging signatures were enriched for CS markers (**Sup Fig2A,B**). This observed cell-type-specific CS enrichment is not explainable by known CS dynamics and was negatively correlated to population-specific proliferation, as determined by the proportion of *Mki67*+ cells (Pearson’s R = -0.24, p = 0.04) (**Sup Fig 2C**). These findings suggest that a universal CS signature may be obfuscated by cellular heterogeneity and that differential expression between subpopulations of cells is not suitable to extract cell-type specific CS markers. Instead, we show that the proportions of cells positive for *Cdkn2a* and other CS markers increase significantly with age in *Tabula Muris Senis* (**Fig 1B**), thus indicating that changes in cell positivity is a more useful metric for identifying dynamic CS genes.

**Figure 1.**
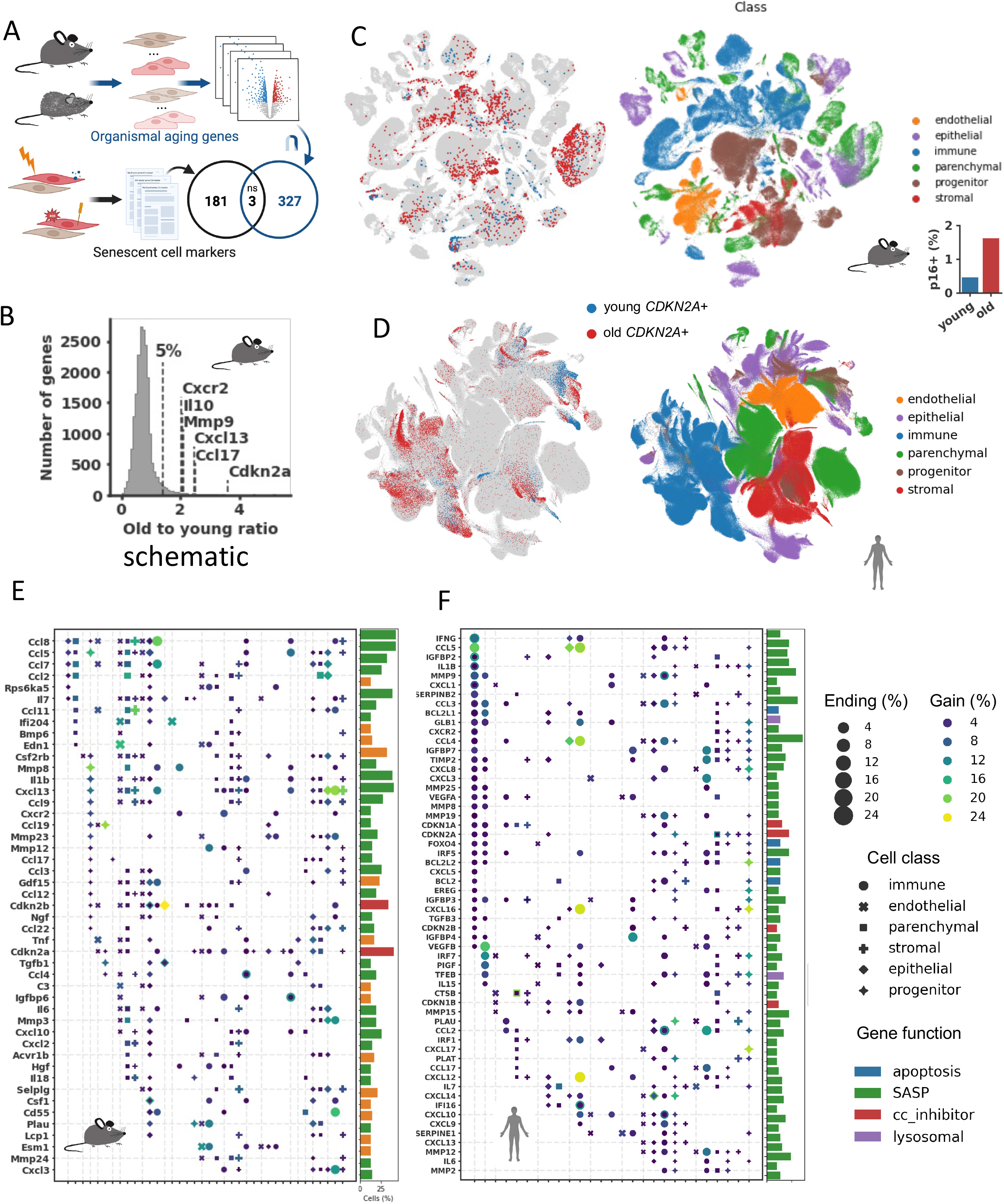
Known senescence markers are cell-type-specific and poorly characterize *in vivo* cellular senescence. (**A**) There is an insignificant overlap between a universal organismal aging signature and previously reported senescence markers (p = 0.58, Hypergeometric). The universal signature are genes present in at least 50% of the cell-specific gene sets elucidated by differential expression between young and old cells previously. (**B**) A histogram depicting the 3-month to 24-month ratio of all cells expressing every gene in the *Tabula Muris Senis* dataset. The top 5% of the distribution is marked by the dashed line and known senescence markers above this are labeled. (**C, D**) UMAPs (left) representing *CDKN2A+* (p16^ink4a^ encoding gene) cells from the mouse and human datasets. Cells from 24-, and 30-month mice are denoted old while 1-, and 3-month mice are young. UMAPs (right) showing all cells in the datasets and are labeled by broad cell classifications. Bar graphs show the percentage of *CDKN2A+* cells relative to all cells in the respective datasets. (**E, F**) Cell-specific maps of marker dynamics in mice and humans. Vertical dashed lines represent the start of a tissue and cell types from that tissue are classified and depicted by shape. Multiple cell types belonging to the same class are overplotted. Gain represents the percent increase of cells expressing the marker relative to young organisms. Bar plots depict the percentage of cell populations which the respective gene is a suitable marker and they are colored by senescence-associated function.

We next analyzed the aging dynamics of CS markers across all tissues and cell types in the mouse and human datasets with respect to the proportion of positive cells. Some of the most widely used markers of CS, such as *Cdkn2a* and *Cxcl13*, showed an overall increase in the proportion of cells expressing these genes with age but had tropism for specific tissue and cell types (**Fig 1C,D, Sup Fig 3A**). Human skin from the face, for example, had appreciable levels of *CDKN2A*+ cells in young individuals and a large increase with age (**Sup Fig 3B**). However, other important markers of CS, such as *CDKN1A* (p21^cip1^ encoding gene), were more constitutively expressed in both young and old mice and humans (**Sup Fig 3C**).

We examined the dynamics of cellular senescence in 60 mouse and 50 human cell types. Overall, the landscape of all CS markers was highly heterogeneous when stratified by tissue and cell type in both species (**Fig 1E,F**). Only 16 of 1,770 pairwise combinations of cell types showed significant senescence marker gene overlap (Hypergeometric, FDR p < 0.05), thus highlighting how senescence program transcription profiles differ widely between tissues and cell types. Cells from the same tissue were most likely to have significant marker overlap (Chi-square, p = 3.2x10^-10^). While fibroblasts from different tissues showed some overlap in senescent cell transcriptional profiles, this was not statistically enriched (Chi-square, p = 0.2). The marker found to increase dynamically with age in the largest number of human cell types was *CCL4*, but this marker was only dynamic in 21 of 50 (42%) cell populations. *Ccl5* and *Ccl8* were the most universally dynamic in mice, but only in 33% of cell types. *CDKN2A* was one of the most enriched markers of senescence in both humans and mice, but it was only dynamic in 26% of human and 32% of mouse cell types. These data indicate that there is no universal senescence marker gene set for all tissues and cell types, and that each cell type within each tissue takes distinct transcriptional paths to the senescence state. Instead, our data maps the suitability of known senescence markers in different organisms, tissues, and cells.

### *De novo* cell-type specific signatures derived algorithmically from single-cell transcriptomes

We used an unbiased computational method to identify putative senescence signatures in distinct mouse and human cell populations (**Fig 2A**). These signatures were derived from 47 mouse and 50 human cell types that spanned wide organismal ages. We leveraged the prior biological knowledge that senescent cells accumulate with increasing organismal age but remain a minority population and this allowed us to identify genes that were dynamic with increasing organismal age within each cell population. Using cell-by-gene positivity matrixes with the selected genes and cells, we constructed networks with hubs of genes likely to be expressed in the same cells. Genes not associated with hubs were removed as they likely represented noise or stochastic processes which increase with organismal age but are not directly related to cellular senescence. Some cell types were comprised of multiple distinct signatures which separated during network analysis (**Fig 2B**). In total, we derived 75 mouse and 65 human putative CS cell-type signatures. The signatures contained both genes as well as values which indicate the degree of correlation they have to other genes within the same signature.

**Figure 2.**
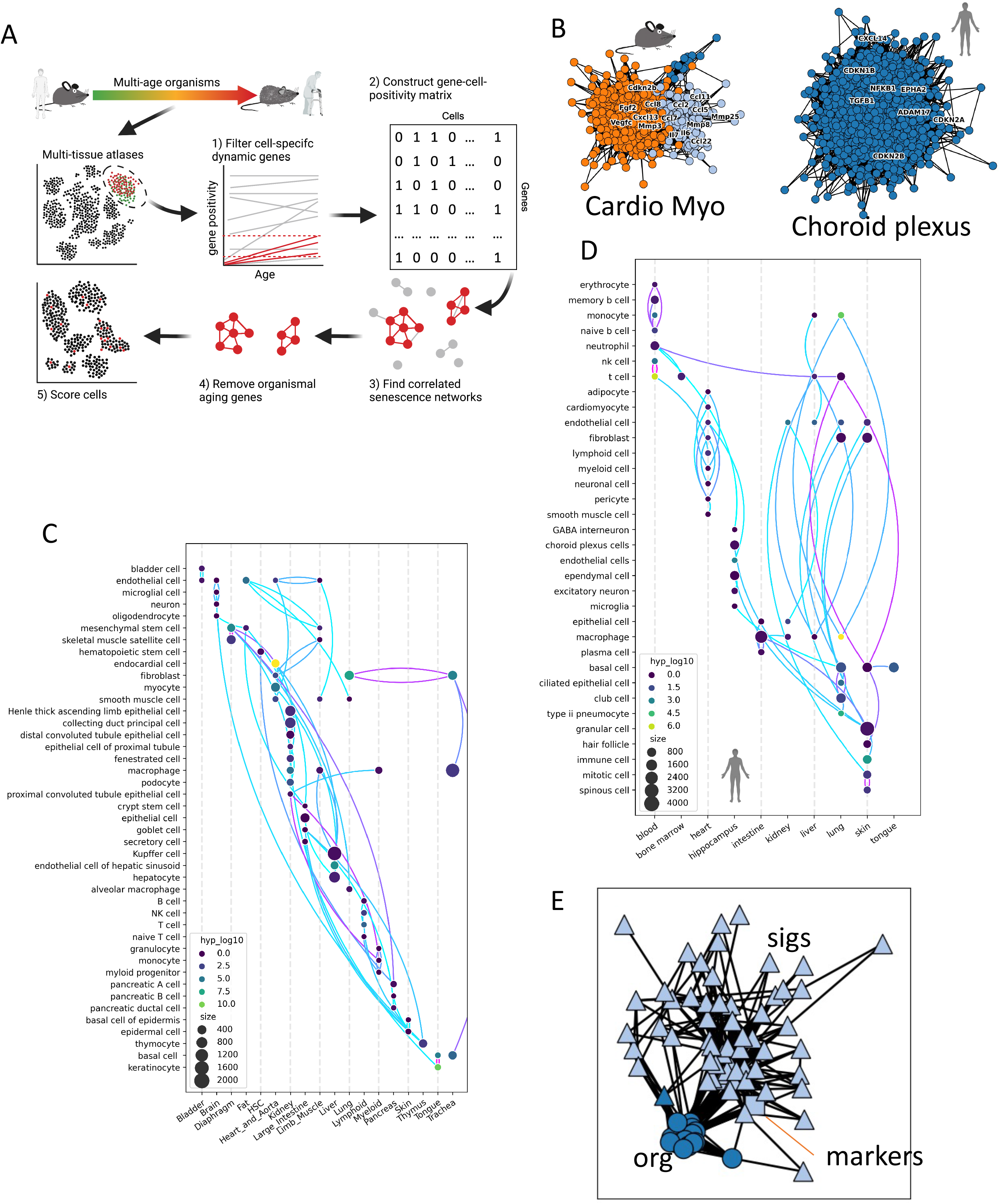
*De novo* cell-type specific signatures derived algorithmically from single-cell transcriptomes. (**A**) Overview of the algorithm used to define cell-specific signatures from mice and humans (see methods). Workflow ends with a cosine signature comparison used to determine signature similarity. (**B**) Example signatures derived from mouse cardiomyocytes and human hippocampal choroid plexus cells. Each node represents a gene and the connections represent positive cell co-positivity. Connections are weighted by Pearson’s R. The colors represent distinct hub signatures within the overall cell signatures. (**C, D**) Representative diagram of all derived signatures from mouse and humans. Each dot represents a signature and is sized by its number of genes. The dot color is the respective enrichment for each signature compared to previously known senescence markers (Hypergeometric). Each signature is connected to its most similar signature and the color of the connection is based on the cosine similarity. (**E**) Network similarity analysis of mouse cell-specific novel senescence signatures and organismal aging signatures. Each shape represents a signature and lines represent significant similarity between them. Similarity (strength of connections) is defined as -log_10_(BH-corrected Hypergeometric P). Network is clustered and colored by Louvain’s algorithm.

Cell-type signatures were highly heterogenous but several known markers of CS were enriched in selected cell types, although not in a consistent manner (**Fig 2C,D**). Importantly, similar gene expression signatures (Hypergeometric FDR < 0.05) were more likely to be found between cells from the same tissue (Chi-square, p = 2.6x10^-9^) than between cells of the same cell type (Chi-square, p = 0.001). However, there were some exceptions in which signatures from cell types found in multiple tissues shared high similarity. For example, senescent fibroblasts found in mouse lungs were most similar to senescent fibroblasts from mouse tracheas and happened to also share the overall highest similarity between any two mouse senescent cell type signatures. However, fibroblast signatures taken from all of the tissues were not more likely to similar to each other (Chi-square, p = 0.3). Overall, the signatures did not form distinct clades when clustered based on similarity, emphasizing heterogeneity and suggesting tissue-specific influences on CS signatures may be more influential than universal drivers of CS (**Sup Fig 4**).

Many cell-type signatures contained overlapping genes despite the high degree of overall signature heterogeneity (**Fig 2E**). There were 387 of 1081 (36%) mouse cell-type-signature pairs that shared significant overlap compared to 16 out of 1770 (0.9%) when using established markers. Based on their genetic profile, cell-type signatures clustered distinctly with each other but not with organismal aging signatures. Importantly, the previously defined set of unspecific senescence markers was more closely associated with the newly derived signatures and fell within the same neighborhood cluster. These observations indicate that the signatures we derived share some underlying genetic characteristics despite being highly distinct and that these signatures likely represent bonified *in vivo* CS programs. Therefore, we developed the open-source *SenePy* Python software package to score single-cells based on their expression of these CS signature genes. *SenePy* rapidly processes thousands of cells and provides relative senescent scores for every cell within a given population.

### Distinct modes and phenotypes of senescence exist within the same cell populations

Cell populations are exposed to multiple stressors and therefore may harbor multiple independent modes of senescence. We observed that multiple cell-type signatures consisted of correlated gene hubs that clustered during network analysis (**Fig 2B, 3A**). Of the 46 mouse cell populations with derived cell-type signatures, 24 contained multiple gene hubs. From 47 human cell populations, 14 signatures were comprised of multiple hubs. Multiple signature hubs likely represent modes of CS with distinct kinetics and gene expression patterns. For example, aging mouse tongue keratinocytes consisted of two hubs, both enriched in established CS marker genes (**Fig 3A**). Gene enrichment analysis indicated different phenotypes and functional roles between cells expressing these separate hubs (**Fig 3B,C**). One of the tongue keratinocyte hubs was more proinflammatory in nature and primarily enriched for immune cell chemotaxis, cytokine, and TNF signaling pathways, typical of the senescence-associated secretory phenotype. We have termed this “type-A” senescence. The other, “type-B”, was enriched for innate immune processes and pathways that typically respond to pathogenic stimuli, for example, the NOD-like receptor signaling pathway (Fisher’s Exact FDR p = 10-8). These observations suggest that these different modes of senescence drive different inflammatory pathways and may contribute differentially to sterile inflammation. Both type-A/B tongue keratinocyte gene hubs were most similar to gene signatures from tongue basal cells, further emphasizing the importance of tissue context on the modes of senescence. However, they were similar to other hubs from multiple tissues and cell types (**Fig 3D**), either presenting as type-A or type-B senescence hubs but not both, suggesting this multimodal pattern is not restricted to tongue keratinocytes. Next, we used *SenePy* to score tongue keratinocytes based on these type-A and type-B signatures. The number of cells that were outliers (> 3σ + µ) in their signature-specific distributions increased significantly as a function of organismal age (**Fig 3E,F**), indicating a higher number of senescent keratinocytes in old mouse tongues compared to young. The increase in senescent cells with age followed a similar pattern in both CS programs but the type-B senescent cells appeared before type-A senescent cells temporally and also reached higher percentages.

**Figure 3.**
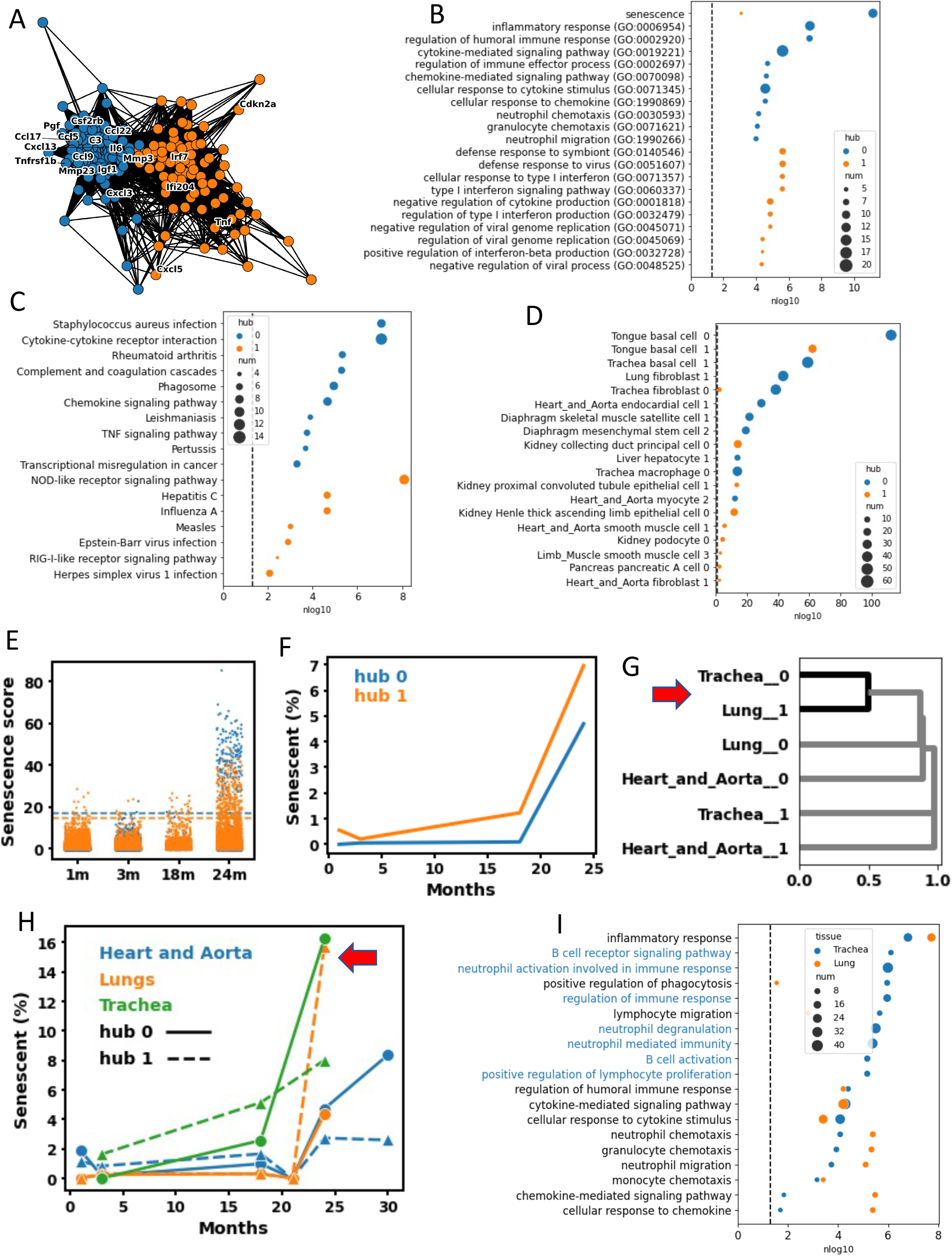
Multiple modes of senescence exist within the same cell populations. (**A**) Signature derived from mouse tongue keratinocytes. Each node represents a gene and the connections represent positive cell co-positivity. Connections are weighted by Pearson’s R. Nodes are colored by Louvain-based assignment to distinct hub signatures. (**B**) GO and (**C**) KEGG gene set enrichment of the two keratinocytes hub signatures. The “senescence” gene set is the pre-defined senescence markers used in this study. Horizonal dashed line represents FDR p = 0.05. (**D**) Pairwise enrichment of the two keratinocyte hubs against all other senescence signatures. (**E**) The strip plot depicts the score of each keratinocyte determined by *SenePy* using the forementioned hubs. Horizontal dashed lines represent three standard deviations above the mean. (**F**) Temporal kinetics of the proportion of cells scored three standard deviations above the mean by *SenePy* for the two keratinocyte hubs. (**G**) Hierarchical clustering of fibroblast hub signatures from mouse lungs, tracheas, and hearts based on cosine similarity. (**H**) Temporal kinetics of the proportion of lung, trachea, and heart fibroblast cells scored high by *SenePy* using their respective signatures. (**I**) GO gene set enrichment of the most similar trachea and lung fibroblast hubs. Gene sets specific to the trachea hub are colored. All enrichment plots use BH-corrected Fisher’s Exact P values.

We next examined senescent fibroblasts from three different tissues to determine the CS characteristics of similar cell types in different contexts. The mouse fibroblasts from hearts, lungs, and tracheas each had two distinct CS signature hubs. Most fibroblast hub signatures shared little genetic similarity and high cosine distance (**Fig 3G**). The senescent cell gene hubs with the highest pairwise similarities were present in the cells of the lungs and trachea, possibly indicative of the spatial proximity and function in the respiratory system may have resulted in similar senescence phenotypes. Functionally, these similar hubs in lungs and trachea shared common biological processes, such as inflammatory response, cytokine signaling, and immune cell chemotaxis (**Fig 3H**). However, the trachea hub was uniquely and highly enriched for genes involved in B-cell signaling, neutrophil activation, and lymphocyte proliferation. When the fibroblast populations were scored with *SenePy*, they showed distinct temporal kinetics (**Fig 3I**). In all cell populations, there was a small proportion of senescent cells in young mice and a drastic increase in old mice. The biggest increase in the proportion of senescent cells occurred between the ages of 18 and 24 months. Interestingly, cells identified using the most similar trachea and lung hubs had comparable temporal kinetics and nearly identical high proportions of senescent cells in 24-month-old mice. Senescent fibroblast populations in the heart and lungs also followed parallel kinetics despite greater gene and ontological distance. Together, these results suggest multiple modes of senescence even within the same populations and that these modes are temporally and phenotypically distinct.

### Cell-specific signatures are unique but share common stress response and inflammatory pathways

We explored the senescence markers we derived for gene and ontological commonality to determine whether there are universal senescence signatures. No gene was present in every signature or hub, but selected genes were statistically overrepresented (random permutations) in the signatures (**Fig 4A**). There were 29 genes with greater than 25% prevalence (n >= 12, p <= 0.0014) in the 47 mouse cell types tested. The most common gene, and the only one above 50% prevalence, was *Hba-a1*, which was present in 28 (60%, p = 5 × 10^−5^) of the 47 signatures. For comparison, *Cdkn2a* (gene that encodes p16) was also overrepresented in the data but only present in 12 signatures (25.5%, p = 0.0014). Among other known senescence markers, multiple Bcl2 family genes, which are common senolytic targets, were overrepresented in these data. Interestingly, along with *Hba-a1*, the hemoglobin subunits *Hba-a2, Hbb-b1*, and *Hbb-b2* were among these most representative genes, despite the absence of erythrocytes in the upstream analysis. Genes present in 8 or more signatures (n = 141, p < 0.01 by random permutation) were enriched for multiple biological processes involved in inflammation, immunity, cytokine signaling, and chemotaxis (**Fig 4B**). The NF-kappa B signaling pathway, a known driver of cell senescence, was among the most commonly enriched pathways (KEGG, BH-corrected p = 0.0004), emphasizing that NF-kappa B plays an important role in some programs of *in vivo* senescence. However, only 9 of the 47 signatures were individually enriched for NF-kappa B signaling, indicating that it is far from universal (**Fib 4C**). Therefore, to test for transcription factors that might be active, we tested our signatures for TF binding enrichment in their gene promoters (**Fig 4D**). The most universally enriched binding motif was RREB1, which was enriched in 38 of 47 signatures. Additionally, we found 46 other transcription factors enriched in over 50% of the mouse signatures.

**Figure 4.**
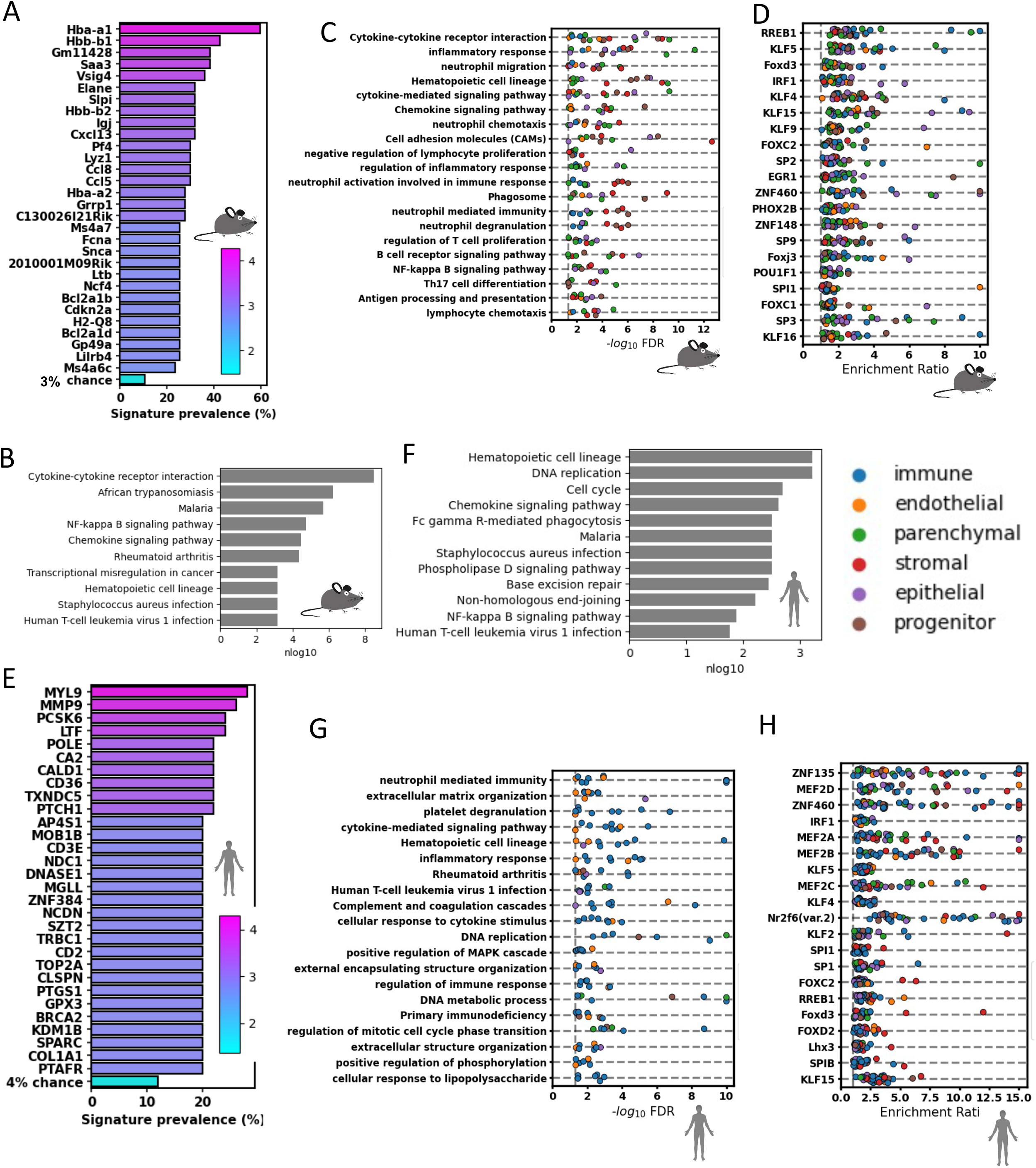
Cell-specific signatures are unique but share some genes and biological pathways. (**A**) Plot depicting the most commonly found genes from the novel mouse cell-specific signatures. Significance was determined from 1000 random sets of signatures equal in size from the background set of genes. (**B**) The 12 most enriched KEGG pathways from a “universal” signature comprised of genes with a high mouse signature prevalence (p < 0.01). (**C**) The most commonly enriched KEGG and GO gene sets in every mouse signature. The bars note the percent of signatures enriched for the given pathway. (**D**) The most commonly enriched transcription factor motifs in the promotors of signature genes from mice. Bars represent the percent of signatures enriched for the given pathway. (**E**) Plots depicting the most commonly found genes from the novel human cell-specific signatures. (**F**) The 12 most enriched KEGG pathways from a “universal” signature comprised of genes with a high human signature prevalence (p < 0.04). (**G**) The most commonly enriched KEGG and GO gene sets in every human signature. The bars note the percent of signatures enriched for the given pathway. (**D**) The most commonly enriched transcription factor motifs in the promotors of signature genes from humans. Bars represent the percent of signatures enriched for the given pathway.

In human cells, only two genes were present in >25% of the human senescence signatures (**Fig 4E**): Myosin light chain 9 (*MYL9*) and matrix metallopeptidase 9 (*MMP9*). *MMP9* is a known SASP component and is present in our curated set of senescence markers. In comparison, *CDKN2A* was only present in 8 of 51 signatures but still higher than what would be expected by random chance (p = 0.01). There were an additional 222 genes present in 8 or more of the cell types. Genes present in more signatures than expected by chance (p < 0.05) were enriched for DNA replication and repair, chemokine signaling, cell cycle inhibition, NF-kappa B signaling, and other biological pathways (**Fig 4F**). When signatures were tested individually, the most commonly enriched pathways included neutrophil-mediated immunity, platelet degranulation, cytokine signaling pathways, and other inflammation and immune pathways (**Fig 4G**), consistent with SASP.

The most universal genes and pathways active in the cell-type signatures from both species shared some characteristics. There were 46 common genes that were enriched in both the mouse and human cell-type signatures. This was only a marginal overrepresentation compared to random chance (Hypergeometric, p = 0.09), suggesting low gene-wise concordance between species. *CDKN2A, CXCR2*, and *CCL3* were the only common genes that were previously established senescence markers. However, the pathway concordance between species was high and both sets of common genes were enriched for 32 common pathways, such as NF-kappa B signaling, AGE-RAGE signaling, and chemokine signaling. Likewise, eight of the top 20 most commonly enriched transcription factors from mouse signatures were also commonly enriched in human signatures (**Fig 4H**). This suggests that core senescence pathways between species are conserved but the individual genes that are enriched in senescent cells show a high degree of genetic variation. Together, these data indicate that our *de novo* cell-type signatures are enriched for known senescence phenotypes and share some commonality between cell types and species despite their high degree of heterogeneity.

### The cell-specific kinetics of senescent cell accumulation with organismal age

We next used *SenePy* to determine the proportion of senescent cells in distinct populations from young and old organisms. The numbers and proportions of cells identified as senescent increased drastically with age in both mice (**Fig 5A**) and humans (**Fig 5B**) overall but showed distinct cellular tropism and kinetics. Within the tested mouse cells, kidney epithelial cells had the highest proportion of senescent cells at old age, followed by tracheal fibroblasts (**Fig 5C**). Out of the 47 tested cell populations in humans with appreciable senescence signatures, type II pneumocytes had the highest percentage of senescent cells reaching a level of greater than 11% in patients aged 68-77. This was followed by several other populations of skin and lung cells (**Fig 5D**). However, in the human cells we observed temporal heterogeneity between patients. For example, cardiomyocytes from a patient in the 58-year age bin were 7.6% senescent while a patient in the 68-year bin had only 0.4% senescent cardiomyocytes. This disparate pattern of fewer senescent heart cells in the older patient persisted in every other heart cell type except adipocytes, suggesting patient-specific differences are important drivers for whole-tissue senescence. In younger individuals (ages 40-60), the heart cell type with the highest proportion of senescent cells was invariably lymphoid cells (**Fig 5E**). Conversely, there was a large increase in the proportion of senescent cells within solid tissue cell types taken from most 60-year+ donors. One 68-year-old patient in particular had a surprisingly high proportion of cardiomyocytes and fibroblasts with high senescence scores. But in the oldest heart tested, there was a relatively low proportion of senescent cells in all cell types except for lymphocytes. These data could be indicative of how circulating cells may be able to promote tissue senescence after transmigrating into tissues or have reduced efficacy in clearing senescent cells.

**Figure 5.**
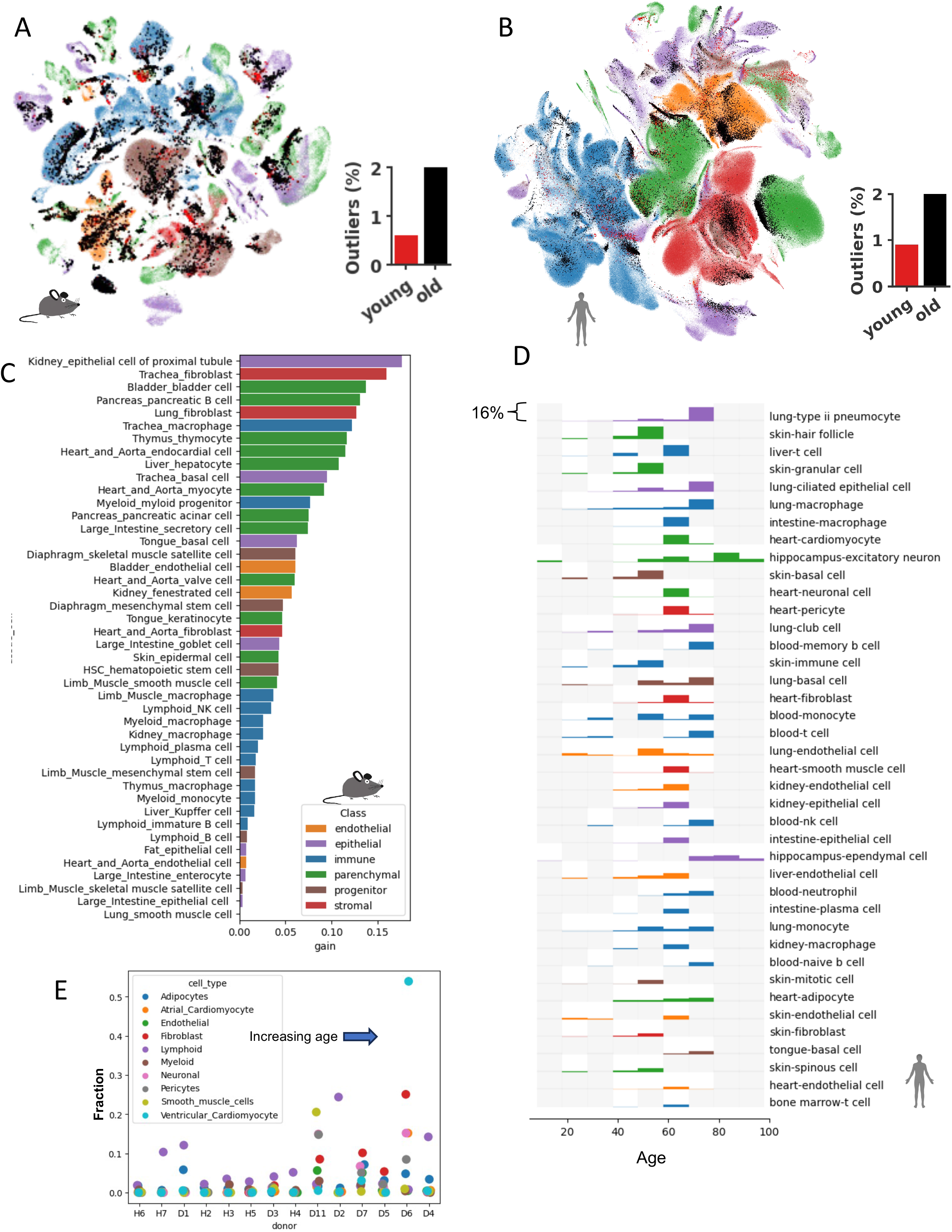
The cell-specific kinetics of senescent cell accumulation with organismal age. UMAPs of (**A**) mouse and (**B**) human cells depicting broad cell classification and overlayed with cells which were outliers determined by their *SenePy* score. (**C**) The proportional increase of *SenePy* outlier cells in old mice (24-or 30-month) relative to 3-month-old mice. (**D**) The proportion of *SenePy* human outlier cells in across age bins. Each row represents 0-16% and grey rectangles note that no data is available. (**E**) The fraction of *SenPy* outliers in individual cell types stratified by heart tissue donor. Age increases along the x-axis from left to right.

The proportions of cells within populations predicted to be senescent by *SenePy* were not correlated to the replicative potential in their respective populations (**Sup Fig 5A,B**). Surprisingly, replicative populations, such as large intestine enterocytes, had minimal increases in the proportion of senescent cells with age. This is corroborated by an undetectable change in the number of *Cdkn2a*+ enterocytes with age and the general lack of correlation between *Ckdn2a*+ cells in replicative populations. Likewise, we did not observe a negative correlation between the population-level expression of telomerase and the calculated senescence burden (**Sup Fig 5C,D**). Thus, indicating that the well-known driver of *in vitro* senescence, telomere attrition, is not the primary driver of senescence within organisms.

### *SenePy* identifies ground-truth *in vivo* senescence more robustly than established markers of senescence

To test *SenePy*’s ability to robustly detect senescent cells in additional datasets, we utilized single-cell RNA-seq data from p16-Cre^ERE2^-tdTomato reporter mice in which p16+ cells become red fluorescent^31^. We first performed differential expression analysis between the tdTomato+ and tdTomato-kidney cells (**Fig 6A**). The mRNAs that were differentially abundant in tdTomato+ cells were enriched for kidney-specific signatures derived in this study. Our kidney-specific signatures were more enriched than previously defined gene sets or a senescence gene panel derived from the AI model GPT-4 that was tasked to identify senescence signatures (n = 100 genes, “senGPT”) (**Fig 6B**). A notable shift was observed in the distribution of cells scored with *SenePy* using kidney-specific signatures but not in a heart endothelial cell hub (**Fig 6C**). Likewise, the mRNAs more abundant in tdTomato+ liver cells were more enriched for *SenePy’s* liver Kupffer cell signature than any other gene sets (**Fig 6D**). These data indicate that *SenePy’*s senescence signatures and derived senescence scores identify cells that become p16+.

**Figure 6.**
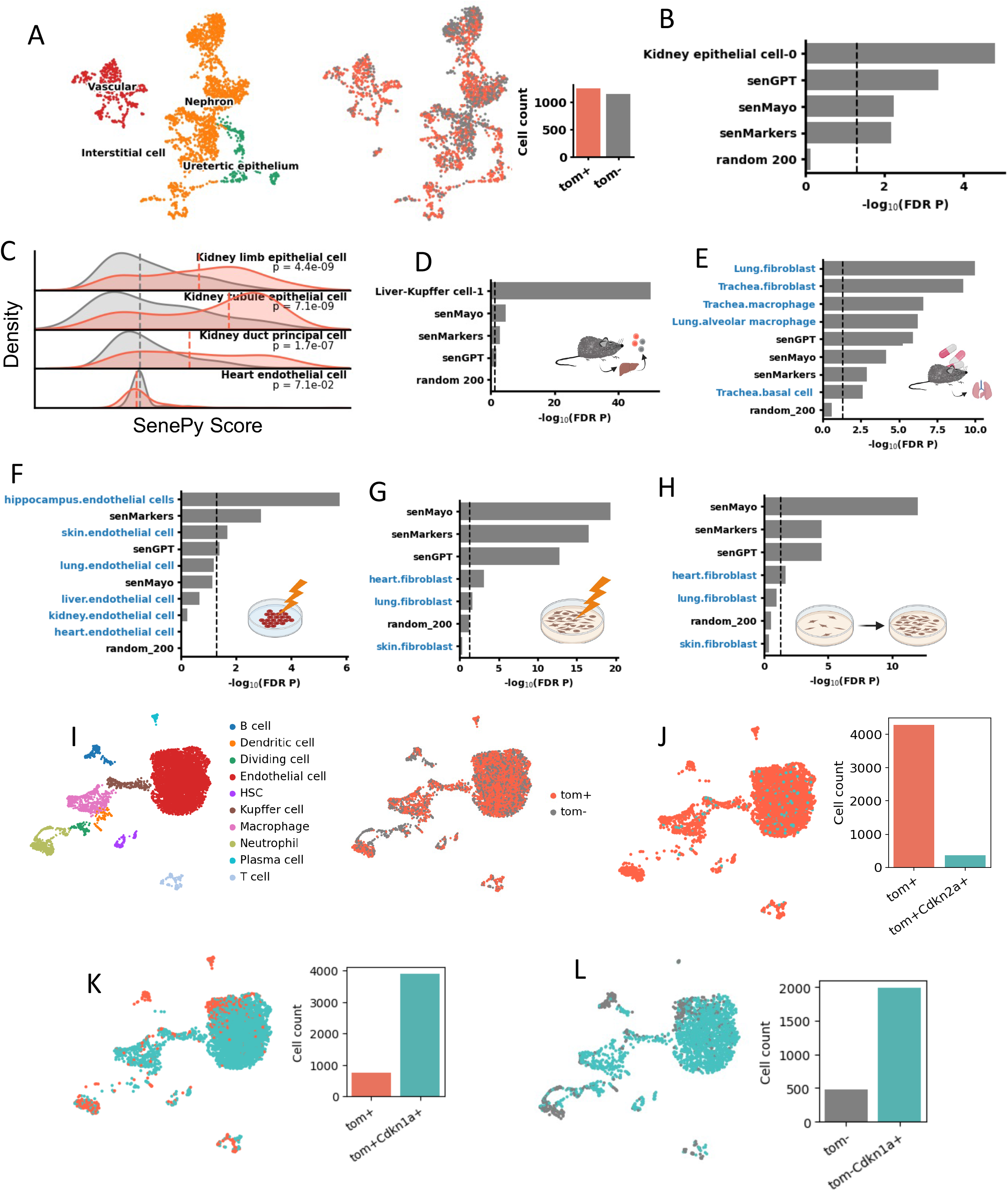
SenePy identifies ground-truth in vivo senescence more robustly than established markers of senescence. (**A**) UMAPs of single-cells from the kidney which were enriched for td-Tomato+ cells. (**B**) Enrichment analysis of differentially abundant genes in the td-Tomato+ kidney cells. Gene sets were derived in this study and we also include the SenMayo signature. (**C**) Density plots depicting *SenePy* score distributions calculated in kidney cells using kidney-specific *SenePy* signatures. (**D**) Enrichment analysis of differentially abundant genes in the td-Tomato+ liver cells. (**E**) Enrichment analysis of lung tissue genes downregulated after senolytic treatment in mice. Blue lables indicate *SenePy* signatures. (**F**) Enrichment analysis of gene mRNAs more abundant in irradiated endothelial cells and (**G**) fibroblasts. (**H**) Enrichment analysis of gene mRNAs more abundant in replicative fibroblast senescence. (**I**) UMAPs of single-cells from the liver which were enriched for td-Tomato+ cells. (**J**) UMAP and representative bar graph of td-Tomato+ cells with *Cdkn2a*+ cells labeled. (**K**) UMAPs and representative bar graphs of td-Tomato+ and (**L**) td-Tomato-cells with *Cdkn1a*+ cells labeled.

We also examined the ability of *SenePy* signatures to identify transcriptional changes due to senolytic treatment as well as those seen in experimental conditions that induce senescence in multiple *in vivo* and *in vitro* models. The genes downregulated in mouse lungs following therapeutic senolysis were enriched for multiple *SenePy* lung-or airway-specific signatures (**Fig 6E**). *SenePy* signatures were more enriched than cell-type agnostic gene sets. *SenePy* does not contain a specific skeletal muscle senescence signature due to data availability, yet mRNA less abundant in mouse muscle tissue following senolytic treatment was enriched for multiple *SenePy* signatures, including one from myocytes (**Sup Fig 6A**). Multiple *SenePy* endothelial signatures were enriched in mRNA more abundant after radiation-induced senescence of human endothelial cells but the advantage of *SenePy* over other gene sets was diminished in this *in vitro* context (**Fig 6F, Sup Fig 6B**). The discrepancy between the *in vitro* and *in vivo* efficacy of *SenePy* was even more apparent in models of senescent fibroblasts in culture (**Fig 6G,H**). In these *in vitro* contexts, *SenePy* was outperformed. These data suggest that *SenePy* recapitulates *in vivo* cellular senescence and that gene sets derived primarily through previous *in vitro* experiments do not.

We next tested the marker suitability of genes that encode for p16^ink4a^ and p21^cip1^ in the p16-Cre^ERE2^-tdTomato liver cells (**Fig 6I**). Only 8% of p16^high^ cells, as indicated by tdTomato, had detectible levels of *Cdkn2a* RNA (p16 encoding gene) (**Fig 6J**). This indicates that either *Cdkn2a* expression was not detectible due to single-cell dropout or the expression of *Cdkn2a* is transitory in the majority of these senescent cells. Another important marker of cellular senescence, *Cdkn1a* (p21 encoding gene), was found in the majority of both tdTomato+ and tdTomato-cells, making its binary expression an inadequate metric because it’s more universally expressed in non-senescent cells (**Fig 6K, L**). These data represent a striking example of why sole reliance on known cellular senescence genes like p16 and p21 is not sufficient, especially in single-cell transcriptomics because of low single-cell resolution, dropout, and marker-independent senescence programs.

### *SenePy* predicts elevated senescence burden in severe disease

We posited that *SenePy* could be applied to datasets to quantify the burden of senescence in disease. We first used *SenePy* to analyze a single-cell RNAseq lung dataset of 19 individuals who died from COVID-19 and 7 age-matched controls^32^ (**Fig 7A**). There were senescent lung cells in both control and COVID-19 patients but there was an observable increase in the senescence of lung epithelial cells among COVID-19 patients (**Fig 7B**). COVID-19 mortality was significantly associated with an increased proportion of senescent lung epithelial cells and were present in AT1, AT2, and general airway epithelial cell populations (Mann-Whitney, p = 0.004) (**Fig 7C**). The proportions of senescence in non-epithelial cells, such as immune cells, fibroblasts, and endothelial cells, were not significantly associated with COVID-19 mortality (**Fig 7D**). Some of the deceased patients had relatively elevated senescence burdens in multiple cell types (**Fig 7E**).

**Figure 7.**
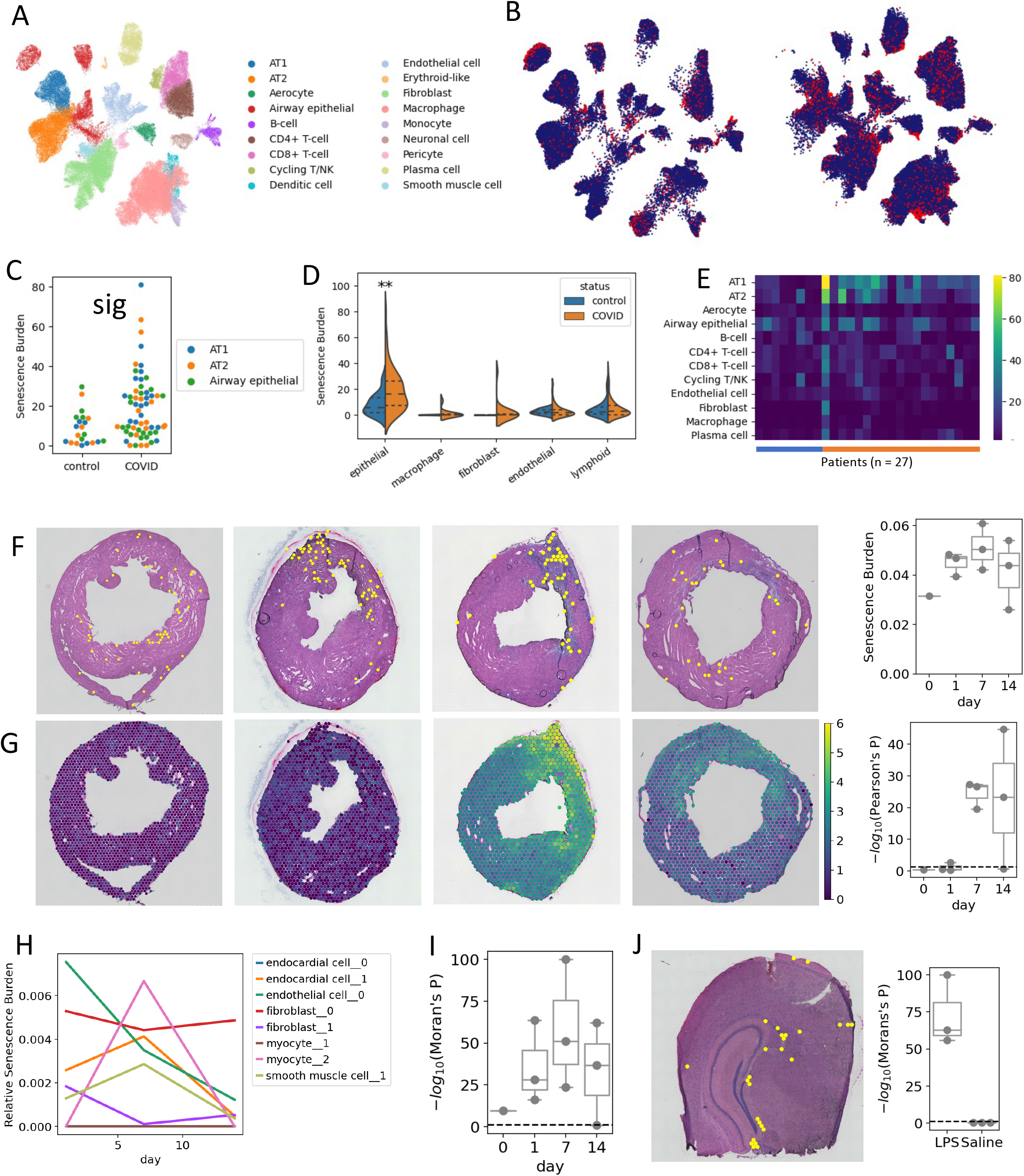
SenePy predicts elevated senescence burden in severe disease. (**A**) UMAP showing cells from uninfected control lungs (n = 7) and patients who died from COVID-19 (n = 22). (**B**) UMAP of control cells (left) and COVID-19 cells (right) with *SenePy* outlier cells labeled. (**C**) Proportion cells identified with *SenePy* from AT1, AT2, and airway epithelial cells (p = 0.004, Mann-Whitney). (**D**) Distributions of *SenePy* score from the major cell lung cell classes. (**E**) Heatmap with the relative senescence burden of each cell type in each patient. (**F**) Representative whole heart H&E staining overlayed with spatially resolved 10x Visium spots. Yellow spots are identified as senescence outliers from their *SenePy* score. Box plot (right) shows the proportion of identified spots at each time point (p = 0.30, one-way ANOVA) (**G**) The bottom images represent post-MI fibrosis via normalized *Col1a1* expression. Scale bars represent 100 µM. The correlation between senescence-like spots and *Col1a1* expression for each sample is shown by the right box plot (-log_10_[Pearson’s R p-value]). (**H**) The relative contribution to the overall calculated senescence burden from the 8 hubs used to score the spots (**I**) Spatial clustering of the senescence-like spots (-log_10_[Moran’s I p-value]). Horizontal line represents p = 0.05. (**J**) Representative H&E image of coronal sections spatially resolved by 10x Visium which were taken from mice 24h after exposure to LPS. Yellow spots are identified as senescence outliers from their *SenePy* score. Summary plot (right) depicts spatial autocorrelation of spots in LPS and saline treated mice (-log_10_[Moran’s I p-value]).

Next, we used *SenePy* to score spatiotemporally resolved mouse transcriptomics data following myocardial infarction^33^. We used the hub signatures we previously derived from mouse hearts to score the spatially resolved spots (**Fig 7F**). Senescent loci were found even in the control heart, corroborating earlier observations that even young organisms have baseline levels of cellular senescence. However, since the spatially resolved spots consist of multiple single cells, the number of single senescent cells in these data are unknowable. The proportion of spots with high senescence burden was highest at day 7 but the change was not statistically significant, likely due to the small sample size (ANOVA, n <= 3). We observed a strong spatial association between spots with high senescence burden and heart fibrosis after MI (**Fig 7G**). Senescent-like spots strongly colocalized in regions of the hearts expressing fibrotic markers such as *Col1a1*. This correlation becomes readily apparent by 7 days post-MI but was not observed in the control heart or hearts shortly after MI. The 8 hubs used to score the spots contributed to the overall senescence burden with distinct temporal patterns (**Fig 7H**). The endothelial cell hub had the highest contribution at day one but continued to decrease up to day 14. The score from a myocyte hub jumped from baseline on day 7 then dropped back down. Other senescence gene programs remained otherwise unperturbed by MI.

Additionally, we observed strong spatial autocorrelation between the spatially-resolved spots with high senescence burden (**Fig 7F, I**). Only one of 9 samples did not have highly significant spatial clustering of senescent-like spots. Unsurprisingly, this same day 14 sample has low senescent spot association to *Col1a1* as well as the lowest overall senescence burden. To further investigate this finding of senescent cell clustering and to see if this phenomenon is apparent in other tissues, we utilized spatial transcriptomics data from mouse brains before and after inflammatory insult (**Fig 7J**). The control brains had a small number of senescent-like spots which prevented significant autocorrelation; however, the brains from LPS-treated animals had amplified senescence signatures and highly significant clustering of senescent-like spots. These data indicate that senescent cells are more likely to be found in close proximity across multiple *in vivo* systems.

## DISCUSSION

There is a paucity of tissue- and cell-specific markers for senescent cells due to the heterogeneity of cells that undergo cellular senescence (CS). This is especially challenging in single-cell transcriptomics because the high rate of dropout and limited sequencing depth of the technology poses a challenge for using classical senescence markers such as *Cdkn2a* and *Cdkn1a* as sole indicators of cell senescence. Furthermore, the cell-specific heterogeneity of cellular senescence represents a major challenge for the development of a universal senescence signature gene panel. Therefore, in this study, we took an unbiased large-data approach to identify cell-specific programs of cellular senescence and created *SenePy* as an open-source platform (https://github.com/jaleesr/SenePy) to identify senescent cells in single-cell transcriptomic data. We validated the *SenePy* approach using single-cell RNA-seq data in p16 reporter mice and applied *SenePy* to determine the kinetics and heterogeneity of senescence across several human and mouse cell types in aging and disease.

Previous studies have generated transcriptomic signatures of cellular senescence based on the generation of senescence in controlled *in vitro* environments ^13, 20^. While these signatures have helped advance the mechanistic understanding of cellular senescence, it is challenging to use such signatures for highly variable *in vivo* contexts. We show that cells express many of these genes at higher rates with age at the organism or tissue level, but none of the genes obtained from such senescence panels were applicable in the majority of cell types tested. Even the widely used marker *CDKN2A* was identified in senescent cells in less than a third of mouse and human cell types. The lack of universality of such senescence marker genes is likely multifactorial, arising from true biological variation and compounded by technical limitations. For example, many SASP genes are also genes that increase with inflammation and organismal aging. A transcript more abundant with age in all tested cells of a population would not be a suitable marker for a phenomenon that is present in only a minority of cells. Furthermore, our analysis of the p16-Cre^ERE2^-tdTomato mouse cells highlights the limits of using a small number of markers such as p16^ink4a^ or *Cdkn2a* in transcriptomics data. While the tdTomato+ cells likely represented bona fide senescent cells^31^, only a small proportion of individual cells had detectible levels of *Cdkn2a* at the time of tissue harvest and sequencing. This likely arose from the transient expression of p16 earlier on in the senescence program in combination with gene dropout inherent in single-cell sequencing. Nevertheless, many highly visible and impactful studies are forced to rely on a small unspecific set of markers because better alternatives did not exist. In addition to finding novel cell-specific signature genes, we provide a cell type suitability map for well-known senescence markers.

Our approach focused on markers specific to individual populations that would not be confounded by transcriptional differences with age. We were able to extract sub-population level programs of CS by setting kinetic thresholds based on the prior knowledge that senescent cells increase with age but are present in the minority of cells and by examining single-cell co-expression as opposed to differential expression. Unsurprisingly, many of our computationally-derived signatures contained and were statistically enriched for pre-established CS markers. However, these comprised the minority and most genes have not been previously thought of as senescence markers. While we did find well-known senescence genes like *CDKN2A, BCL2-*family genes, and various SASP factors to be statistically common within our signatures, the most universal markers were novel ones. For example, in mice, the most common signature gene we identified was the alpha hemoglobin subunit. Hemoglobin has been previously reported as a response gene to oxidative stress in non-erythrocytes^34, 35^, but has yet to be reported in CS. Interestingly, in human signatures there was no significant overrepresentation of globin genes. We observed this and other important differences between mouse and human signatures, suggesting organism-specific CS marker panels to be more specific. Additionally, we used our signatures to map the kinetics of senescent cell accumulation in many different tissues and cell types. Recent work has mapped the abundance of senescent marker mRNA in 13 different tissues as a function of mouse age and in a progeria model^36^. This work, however, was agnostic of cell type and relied on a small set of senescence markers. To our knowledge, this study is the first that comprehensively maps the increase in senescent cells in many different mouse and human tissues with respect to cell type.

The CS marker genes identified in this study may contain novel and cell-specific targets for senolytics. Senolytics have been effective in treating age-associated disease and at extending healthspan in model organisms^7-10^. Many senolytics inhibit anti-apoptotic factors such as BCL2-family proteins^37^, and we indeed found numerous BCL2-family protein genes as markers in many of our cell-specific signatures. Therefore, this indicates that our methodology was able to extract actual senolytic targets. Our data will also inform which tissues and cell types BCL2-family inhibitors and other senolytics are likely more effective. Recent work has used a computational approach to identify a novel surface marker on senescent cells to direct targeted CAR-T senolysis^38^, expanding the potential field of senolytic targets and emphasizing the importance of the novel markers found in our study. Furthermore, clearance of senescent cells is not beneficial in every context and cell-specific targeting of senescent cells may be warranted in some scenarios. For example, liver regeneration is dependent on hepatic stellate cell senescence but is hindered by senescence of the liver parenchyma^39^. Our work uncovers potential cell-specific targets for specific senolytic therapy.

Senescent cells contribute to cardiovascular pathology^6, 40^, but their role in disease has never been spatiotemporally characterized. We show that senescent cells localize at sites of heart fibrosis. The proportion of senescent cells in each heart throughout the time series did not significantly change and were present even in the control heart. We also observed highly significant spatial clustering of senescent foci in the hearts and brains of mice which may be supporting evidence for an *in vivo* bystander effect^15^. The spatial distributions of senescent cells have been previously examined from spatially resolved transcriptomics data in aged mouse brains ^41^. Their results show that *Cdkn2a+* spots are adjacent to activated microglia but no spatial clustering of *Cdkn2a+* spots. Though, their methodology relies on a narrow definition of CS which may not translate to actual p16^ink4a^ and does not account for Cdkn2a dropout or p16-independent forms of senescence. These and other data would greatly benefit from a reexamination with more comprehensive gene sets, such as those proposed herein.

Here we report that senescence burden in human lung endothelium is associated with COVID-19 mortality. Senolytic treatment prior to infection with betacoronavirus has been shown to reduce mortality in mice suggesting that pre-existing senescent cells increase disease risk^9^. In humans, deceased patients infected with SARS-CoV-2 had an increased number of mucosal cells expressing senescence markers and infection itself can induce senescence^4^. Their transcriptomics data also showed CS marker abundance to be highest in epithelial cells relative to other mucosal cell types. Taken together, these studies support our findings that senescent epithelial cells contribute to COVID-19 mortality.

By design, our methodology removes genes that are constitutively expressed at baseline or in aged cells to maintain a distinction from organismal aging. Inherently, this discounts genes that may be part of the senescence program which overlap with the transcriptional shift with age. We also do not account for genes that are downregulated in senescent cells. Negative markers of senescence would add extra information to better identify senescent cells, but to find negative correlations in all pairwise combinations of genes with this methodology was computationally limiting. The comprehensiveness of our signature panel is also limited by the data available at the time of study and the exact set of tissues and cells tested influences any conclusions of universality or comparison between species. We do not expect our signatures to negate the need for large scale future efforts such as SenNet^21^. We instead expect our work to be complementary and assist these efforts.

This work comprehensively identified gene expression programs and signatures of senescent cells that are stratified by species, tissue, and cell type and used them to broadly characterize senescent cells in mice and humans. We created *SenePy*: a computational platform that assigns a senescence score to individual cells in single-cell transcriptomic data, which can serve as a resource to uncover cell-type and tissue-specific mechanisms of cellular senescence *in vivo*.

## METHODS

### Data collection

Single-cell RNA mouse data were collected from the *Tabula Muris Senis* atlas (ref). *Tabula Muris Senis* consists of single-cells from 30 mice from 1 to 30 months of age taken from 19 tissues. Human single-cell data were collected from 7 studies. The liver data were obtained from five donors ranging from 21 to 65 years old^29^. Single skin cells were obtained from 6 patients ranging in age from 18 to 48 years old^25^. Lung data were collected from 17 donors ranging from 21 to 72 years old^28^. Human heart cells were taken from 14 patients ranging from 40 to 75 years old^26^. Human hippocampal cells were collected from 37 patients ranging from newborn to 92 years old^23^. These tissue-specific datasets were given priority for downstream analysis in their respective tissues, but we used additional multi-tissue atlases. Cells from the *Human Cell Landscape* came from 51 donors ranging from 21 to 66 years old and from 25 different tissues^27^. Cells from *Tabula Sapiens* came from 15 patients ranging from 22 to 74 years of age^24^.

### Data annotation

The data were available in a range of formats from fastq to processed and annotated count data. Data from *Tabula Muris Senis, Tabula Sapiens*, and the human heart study were provided with cell type annotations. The Lung and hippocampal studies provided unannotated counts. Fastq data were processed through 10x CellRanger or the Dropseq protocol (https://mccarrolllab.org/dropseq/) depending on the technology used to prepare the libraries. All fastqs were aligned to GRCh38. Processed single-cell counts were handled with Scanpy^42^. Cells were filtered out if they had a relatively low or high number of detected genes or high relative proportion of mitochondrial reads (thresholds varied based on dataset distribution). We used a variety of methods to annotate cell types. Since the *Human Cell Landscape* contained many tissues and cell types, we transferred the annotations from *Tabula Sapiens* using scANVI^43^ after processing the raw data with scVI-tools^44^. If a cluster, which was called by the Leiden algorithm on the scVI embeddings, had lower than 85% cell-type agreement, those cells were not used in downstream analysis. Cell types from the liver, skin, and lung studies were annotated similarly but clusters with poor label transfer were instead manually annotated using known cell-type markers^45^. For lack of a reference dataset, the hippocampal data were annotated exclusively using known markers. Annotations were harmonized across datasets (e.g., “kidney endothelial cell” changed to “endothelial cell”) and mapped back on to the raw counts. Cells lacking annotations because they failed QC or label transfer were discarded. For total dataset visualizations, the species-specific raw data were integrated using scVI and the embeddings were projected via UMAP.

### Mouse cell-type specific age-dynamic genes

Mouse data came from mice aged 1, 3, 18, 21, 24, 30 months (m) but age availability varied by tissue. Cells were stratified by tissue, age, and cell type. The starting baseline was chosen as 3m if there were at least 200 3m cells, if not the starting baseline was aggregated with 1m cells. Likewise, 30m was prioritized for old cells if at least 200 were present, otherwise the old baseline fell back to 24m. The proportion of cells expressing one or more UMI copies of a gene was determined in each population (**Equation 1**). Zero values at 3m or 1m were imputed with the inverse of the cell count (*p*_*age*_ = *n*_*totalcells*_^−1^). Young cells were used as baselines and the ratios in old cells were determined relative to them (**Equation 2**). Genes were only considered dynamic if they were found in less than 5% of young cells (*p*_3*m*¼1*m*_ *j* 0.05), found in less than 20% of old cells (*p*_24*m*0;|30*m*_ *j* 0.2), were in greater than 1% of old cells (*p*_24*m*0;|30*m*_ > 0.01), increased at least 0.5% (*p*_24*m*0;|30*m* -_ − *p*_3*m*0;|1*m*_ > 0.005), and the old to young ratio was greater than 2.5 or the gain was greater than 5% (*r*_24*m*0;|30*m* -_ > 2.5 | *p*_24*m*0;|30*m*_ − *p*_3*m*0;|1*m*_ > 0.05). These thresholds assume that the proportions of cells expressing senescence marker increase with age but remain in the minority. Dynamic genes are cell-specific markers and do not account for small changes to baseline levels of constitutively expressed genes, which may be senescence-associated genes but are not specific markers.

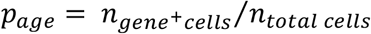

(**Equation 1**) *p*_*age*_: the proportion of cells positive for a gene at a given age. Where *p*_*age*_ *= n*_*gene*+*cells*_ is the number of cells positive for a given gene and *n*_*totalcells*_ is the number of total cells in the same population.

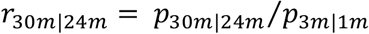

(**Equation 2**) *r*_30*m*0;|24*m*_: ratio of old cells positive for a gene relative to young cells. Where *p*_30*m*0;|24*m*_ represents the proportion of cells positive for a gene in cells from 30-or 24-month-old mice and *p*_3*m*0;|1*m*-_ are the proportion of cells positive for the gene from 3-or 1-month mice.

### Human cell-type specific age-dynamic genes

Human ages were binned into 10-year bins to account for the continuous range of human ages. Bins ranged from 8 (8-17 years old) to 88 (88-97 years old). To be considered for further analysis a cell-type population must 1) have three unique bins with at least 100 cells in each bin or a bin <=28 and a bin >=58 with at least 100 cells and 2) have a bin >=48 with at least 100 cells. These criteria were required in individual datasets because downstream analysis avoids confounding effect from multiple studies. Cells were stratified by dataset, tissue, age bin, and cell type. The proportions of cells expressing genes were calculated for each age bin (**Equation 1**). The young starting populations were selected from the 8, 18, or 28 10-year bins if one was present, else the starting proportion was calculated by regressing the age and known proportion values and solving for 18 years (**Equation 3**). The old ending populations were selected from the oldest age bin (*p*_*old*_). Genes were only considered dynamic if their age by proportion slope was positive (**Equation 4**), their maximum positivity was less than 25% (*p*_*m*ax_ < 0.25), their starting proportion was less than 5% (*p*_*young*_ < 0.05), their gain was greater than 1.5% (*p*_*old*_ − *p*_*young*_ > 0.015), the maximum proportion was not two times higher than the ending proportion (2*p*_*m*ax_ < *p*_*old*_), and the maximum proportion had to be in the 48 year bin or higher (max(*p*_48+_) *= p*_max_).

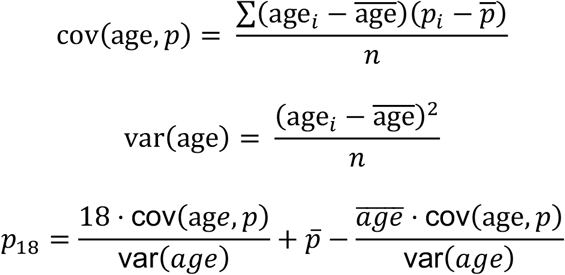

(**Equation 3**) *p*_18_: extrapolated proportion of cells positive in the 18-year bin. Where cov(*age, p*) represents the covariance between age and proportion *p*, var(*age*) is the variance of age, and *n* is the sample size.

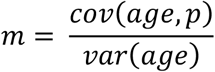

(**Equation 4**) *m*: slope of the linear regression line for the proportions of a given gene with age. See equation 3.

### Identifying novel senescence signatures from mice

Age-dynamic genes for each tissue-cell-type were found as described earlier. Count data were subset by these genes and further subset to only include cells from mice >21m. Subsets with fewer than 100 cells were not tested further. The count matrixes were binarized to represent cell by gene positivity. Every pairwise combination of genes was tested for Pearson’s correlation (**Equation 5**). To test the statistical significance, each pairwise comparison was randomly permutated 500 times. Pairwise correlations were kept if they had a positive r value and if their r value was at least 0.05 higher than the respective q99 r value from the random permutations (**Equation 6**). The filtered correlations were used to construct networks with NetworkX (https://networkx.org/). The Louvain algorithm was used to group genes into clusters. Network clusters with fewer than 5 genes or genes with no correlations were removed. Genes loosely connected to clusters were removed if they had fewer than log(*n*_*cluster genes*_) (Where *n*_*cluster genes*_ is the number of genes in a Louvain cluster) connections to other genes in the network. The cleaned clusters are hereby referred to as hubs and the aggregated hubs for each cell type as a cell-specific signatures.

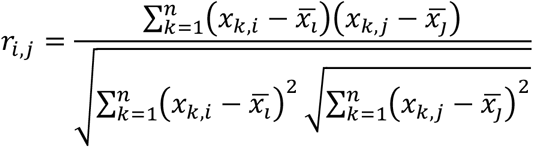

(**Equation 5**) *r*_*i,j*_: Pearson’s correlation coefficent for dynamic genes *i* and *j*. Where *x*_*k,i*_ represents the binary expression value of gene *i* in cell *k*; *x*_*k,j*_ is the binary expression value of gene *j* in cell *k*; and n is the total number of cells in the population.

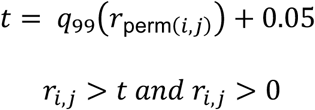

(**Equation 6**) *t*: significance threshold. Where *r*_perm(*i,j*)_ represents the distribution of correlation coefficients for 500 random permutations of gene *i* and *k. q*_99_ represents the 99^th^ percentile value of this distribution. The inequality depicts one criteria of gene selection based on *t*.

### Identifying novel senescence signatures from humans

Age-dynamic genes for each dataset-tissue-cell-type were found as described earlier. Count data were then subset by these genes and further subset to only include cells from patients 48 years of age or older. Significant correlations, networks, hubs, and signatures were generated similarly to those from mice.

### Novel signature comparison

Each signature or hub has a set of genes and corresponding weights for how many connections a gene shares with other genes. Pairwise cosine similarity was calculated by comparing the union of each gene list and imputing 0s (**Equation 7**).For pairwise hypergeometric similarity between two signatures, the cumulative distribution function for two lists of genes was determined using the genes present in the original species aggregated counts as the background list (**Equation 8**). For signature network analysis, all pairwise hypergeometric sf values (i.e., p-values) were corrected with the Bonferroni method, converted to − log_10_ *sf*_*corrected*_, and used as similarity scores between signatures if they were significant.

To find genes represented in the signatures more than expected by chance, we used a random permutation method. A set of hubs with random genes identical in size to the original signatures were generated 1000 times from the background set of expressed genes in the dataset. A distribution was created representing the number of times a gene was found in each of the 1000 permutations. The actual number of signatures a gene was found in was compared to this distribution to determine significance.

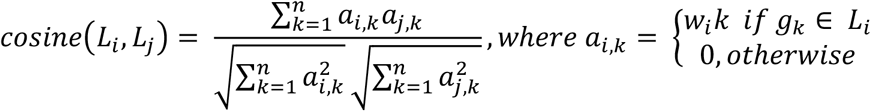

(**Equation 7**) *cosine*(*L*_*i*_, *L*_*j*_): cosine distance between two signatures *L*_*i*_ and *L*_*j*_. Where *a*_*i,k*_ represent the weight of gene *k* in gene list *i*, and is equal to the corresponding connection value *w*_*i*_*k* if gene *g*_*k*_ is in gene list *i*, or 0 otherwise. Where n represents the total number of genes in the union of signatures *L*_*i*_ and *L*_*j*_.

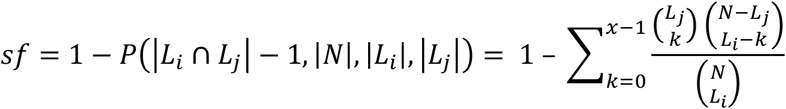

(**Equation 8**) *sf*: survival function of the hypergeometric distribution *P*(*x, N, I, j*). Where *L*_*i*_ and *L*_*j*_ represent two gene lists and *N* represents the background gene list. *x* is equal to the cardinality of the *L*_*i*_ and *L*_*j*_ intersection minus 1.

### Gene set enrichment – GO, KEGG, transcription factor binding

We used the Enrichr python API gseapy^46^ for gene set enrichment against the GO and KEGG databases (refs). The background set of genes used came from all expressed genes from their respective datasets. Only FDR-corrected p-values below 0.05 were considered significant. A custom “senescence” gene set was added which was comprised of the union between all literature-based senescence markers collected for this study and senMayo^20^.

For transcription factor binding analysis, the regions 1000 bp upstream and 500 bp downstream of the transcription start sites were extracted for each gene in a gene list. JASPAR 2020 core vertebrate non-redundant position frequency matrices were us as the input motifs (ref). The extracted regions were examined for relative motif enrichment using the MEME-suite simple enrichment analysis.

Only Benjamini-Hochberg-corrected p-values below 0.05 were considered significant.

### Scoring cells using SenePy

Gene signatures are comprised of genes and their respective number of edges in their network (termed importance value). We developed SenePy, a lightweight and fast scoring algorithm specific for our gene sets that borrows from Seurat’s AddModuleScore() and Scanpy’s tl.score_genes(). SenePY is built in python and integrates well with scanpy and anndata. SenePy has four core functions: load_hubs(), translator(), score_hub(), and score_all_cells(). The load_hubs() function initializes the hub object which includes the hubs themselves along with additional metadata, such as each hub’s enrichment for known senescence genes. Depending on the input data and its respective reference, the optional translator() function can be used to harmonize gene symbols based on known gene aliases. The score_hub() function takes one input hub and anndata and returns a list of scores for each cell. The score_all_cells() takes one input hub and anndata and stratifies the data based on input categories, for example, to score individual cell types separately to avoid confounding the score.

The scoring happens in multiple steps. First, the mean is calculated for each gene in the dataset across all cells (**Equation 9**). All genes are ranked by their mean and split into n_bins_ (default: 25) expression bins (**Equation 10**). Next, n_ctrl_size_ (default: 50) background genes are selected for each input signature gene from its corresponding expression bin (**Equation 11**). The counts data are then optionally binarized (default: True) to represent the binary senescence cellular state and the gene-cell positivity from which the underlying networks were derived . Next, the counts are optionally amplified (Default: True) by their corresponding importance value from the input signature (e.g., if [*Cdkn2a*, 2] is in the signature all *Cdkn2a* values would be multiplied by 2) (**Equation 12**). Then the cell-by-signature-gene matrix is averaged across the cell axis and subtracted from the mean of the cell-by-background matrix also averaged across the cell axis (**Equation 13**).

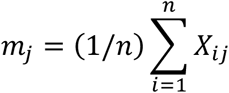

(**Equation 9**) Where *X* is a matrix which contains the expression level of gene *j* in cell *i* and *m*_*j*_ is the average expression of gene *j* across all cells.

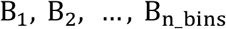

(**Equation 10**) Where *B*_*n_bins*_ is the number of bins used to categorize every gene based on their mean expression.

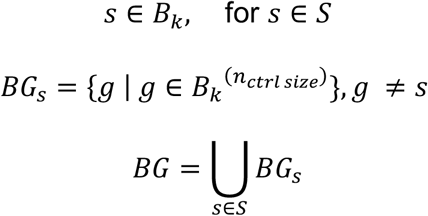

(**Equation 11**) *BG*: background gene set. Where *S* is the gene signature and *s* is a gene within *S. B*_*k*_ is a subset of genes that fall within the k-th expression bin based on their mean expression. Where *g* is a background gene selected from expression bin *B*_*k*_ and *n*_*ctrl size*_ is the number of background genes selected for each signature gene *s* from the corresponding expression bin. Where *BG*_*s*_ is a set of background genes randomly selected from the same expression bin *B*_*k*_ as the signature gene *s. BG* is the union of all background genes selected for each signature gene *s*.

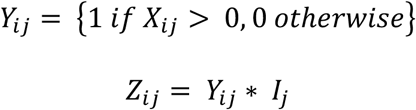

(**Equation 12**) *Z*_*ij*_: modified expression matrix. Where *Y*_*ij*_ is the optionally binarized expression matrix *X*_*ij*_ and *I*_*j*_ represents the optional importance values for gene *j*.

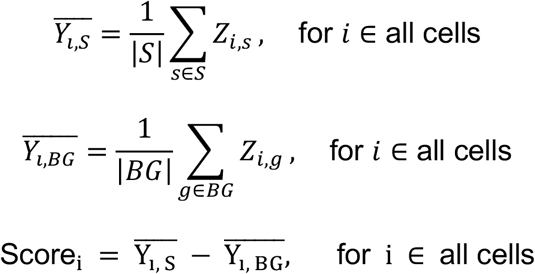

(**Equation 13**) Score_*i*_: *SenePy* score for cell *i*. Where |*S*| and |*BG*| are the cardinality (number of elements in the set) of gene signatures *S* and *BG*. Where *Z*_*i*,s_ and *Z*_*i*,g_ represent the optionally amplified expression values of the genes in the gene signature *S* and background gene set *BG*, respectively.

### Senescence burden in spatially resolved transcriptomics

Data was preprocessed in Scanpy and pots with fewer than 1000 detected genes were removed. Cells were normalized to 10,000 counts and log converted. The 8 heart-specific mouse hub signatures were used to score the spatially resolved mouse hearts independently using senepy.score_hub() with a translator() and with binarize and importance set to False because Visium data has higher gene counts than single-cell data. Outlier spots were identified in each sample if they fell 3 standard deviations outside the mean for their respective sample distribution in addition to a combined sample distribution (*Outlier* > *µ* + 3*σ*). The outliers from each signature were merged to determine if any given spot was an outlier. Relative senescence burden is presented as the proportion of outlier spots. For the mouse brains, we used the top 150 most common genes in all the signatures because we had no specific mouse brain signatures. To determine spatial autocorrelation, we used the ESDA python package (https://pysal.org/esda/). The weights of the autocorrelation were weighted by the inverse of the Euclidean distance between two spots with a value of 1 to denote an outlier and 0 for normal spots. Three is used as a maximum value for Euclidean distance and the weights for distances beyond three are set to 0 (**Equation 14**).

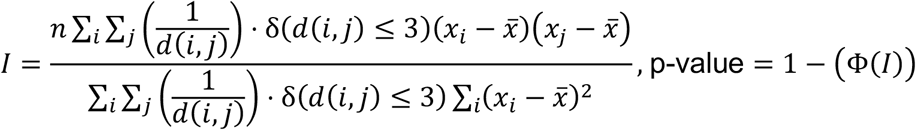

(**Equation 14**) *I*: Moran’s I. Where *n* is the number of spots; *d*(*i, j*) is the Euclidean distance between spot *i* and spot *j*; *δ*(*d*(*i, j*) ≤ 3) is 1 if the distance is greater or equal to 3 and otherwise 0; and *x*_*i*_ and *x*_*j*_ are the values at spot *i* and spot *j*. Φ(*I*) is the CDF of the standard normal distribution at the Moran’s I value.

### Senescence burden in COVID-19 mortality

Single-cell lung data from 20 patients that died from COVID-19 and 7 control patients were collected from an available atlas^32^. Doublets were removed from each individual sample using SOLO^47^ in combination with SCVI-tools. Cells with low counts or high mitochondrial reads were removed. SCVI-tools was used to integrate the 27 samples, using sample ID as a categorical covariate and mitochondrial read percent, ribosomal read percent, and total counts as continuous covariates. Cell types were manually annotated using known cell-type markers (PanglaoDB). Cell types were scored with respective cell type hubs from SenePY (e.g., epithelial cells were scored with ciliated epithelial, basal cell, club cell, and pneumocyte hubs) using the senepy.score_all_cells() function. Cells were divided and scored as individual subtypes (e.g., AT1, AT2, airway epithelium). Cell outliers were identified in each sample if they fell 3 standard deviations outside the mean within every respective cell-type distribution (*Outlier* > *µ* + 3*σ*). Outliers were merged across hubs to identify all cells with potential senescence burden and output as a proportion of total cells.

---

**Supplemental figure 1.**
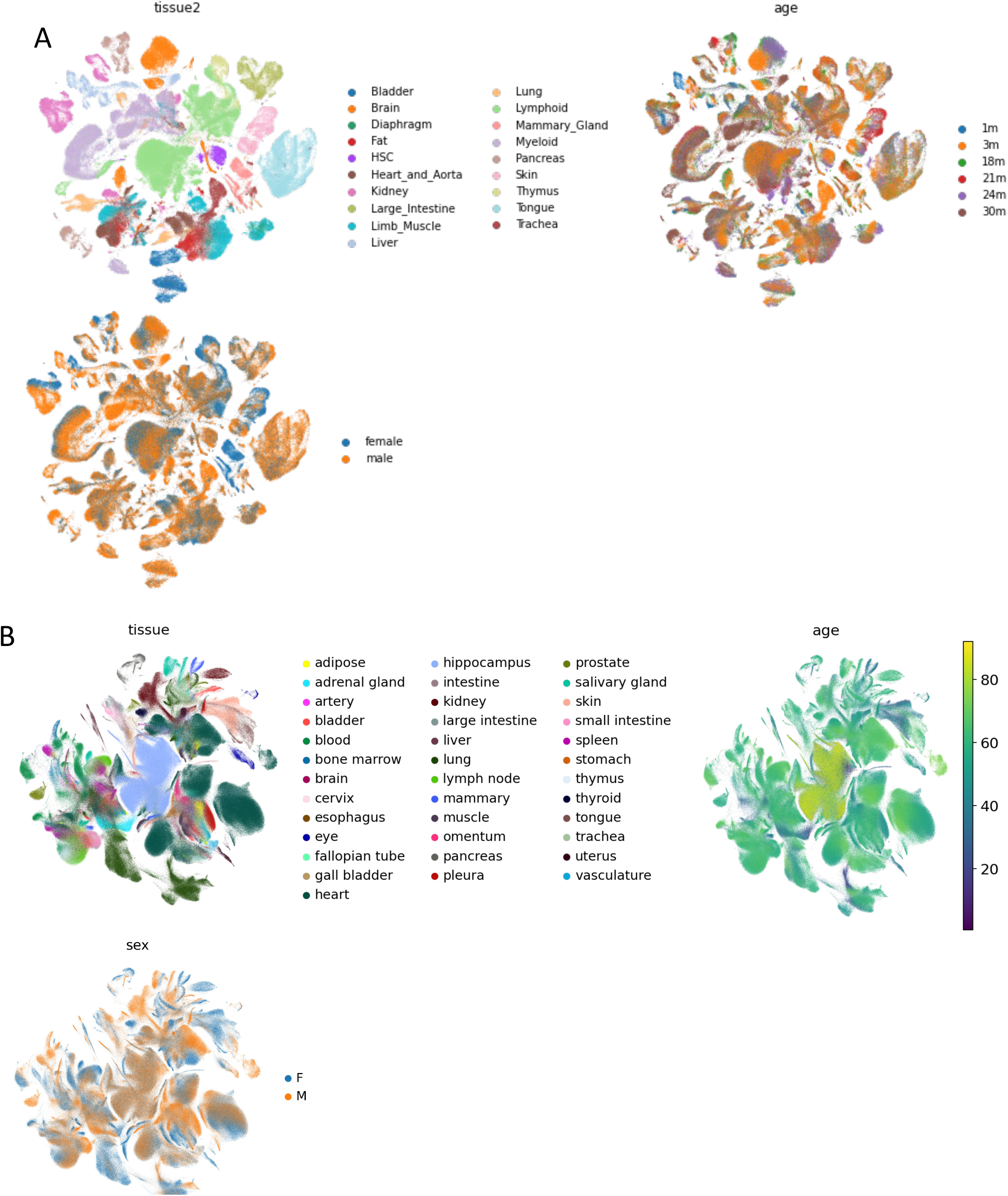
(**A, B**) UMAPs of all mouse and human cells used that are colored by tissue of origin, age, and sex.

**Supplemental figure 2.**
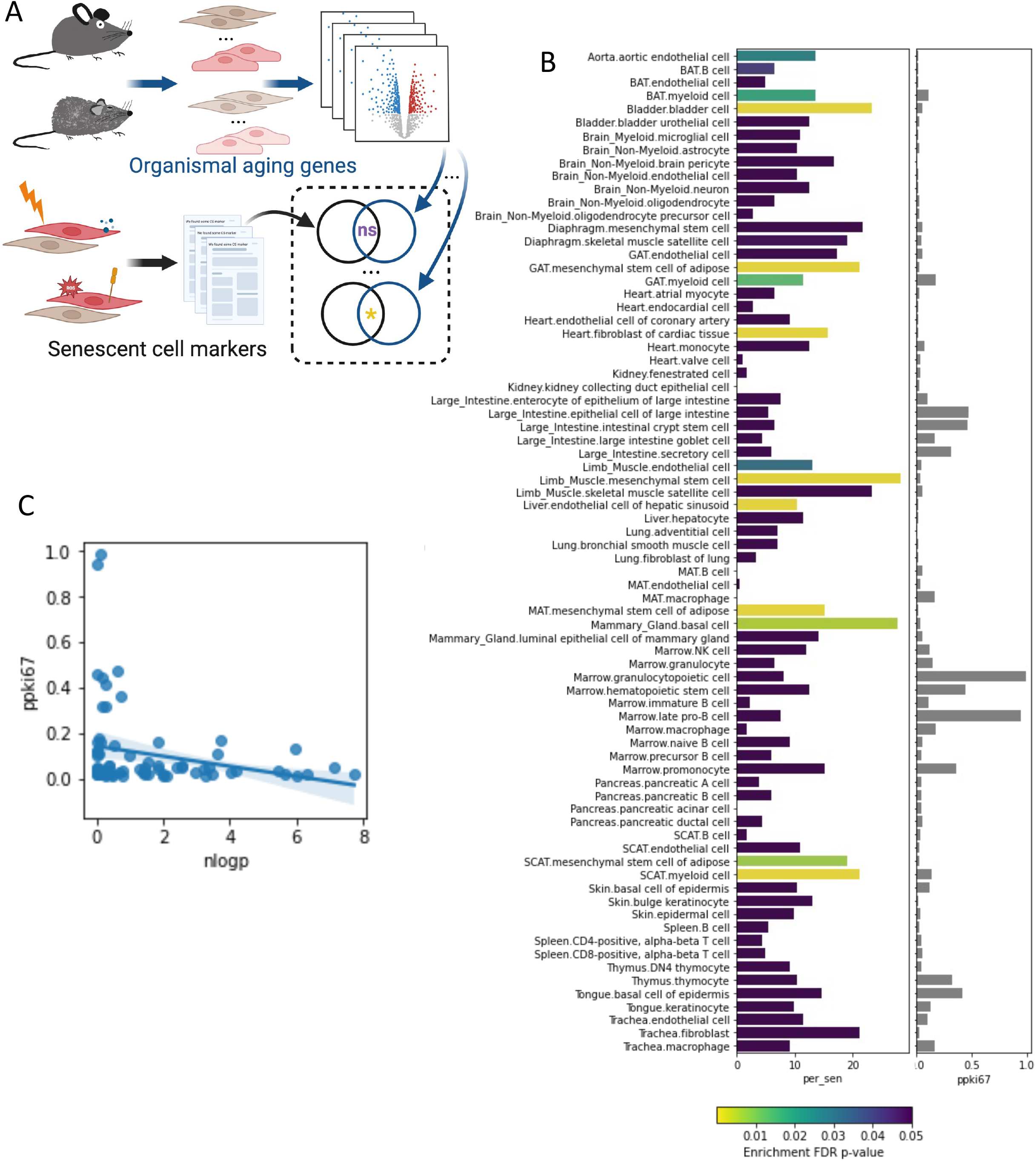
(**A**) Overview of the senescence marker gene set enrichment analysis. Cell-specific organismal aging signatures determined in a previous study were compared to a curated set of senescence markers taken from multiple studies. (**B**) The statistical enrichment of every organismal aging signature for senescence markers. Color represents the significance of enrichment (Hypergeometric, BH-corrected) and the size of the bar represents the percentage of senescence markers present in the aging signature. The aligned plot on the right shows the proportion of cells in each population that were expressing the proliferation marker Ki-67. (**C**) There is a negative correlation between senescence gene set enrichment and the proliferation state of a population (p = 0.038, Pearson’s R).

**Supplemental Figure 3.**
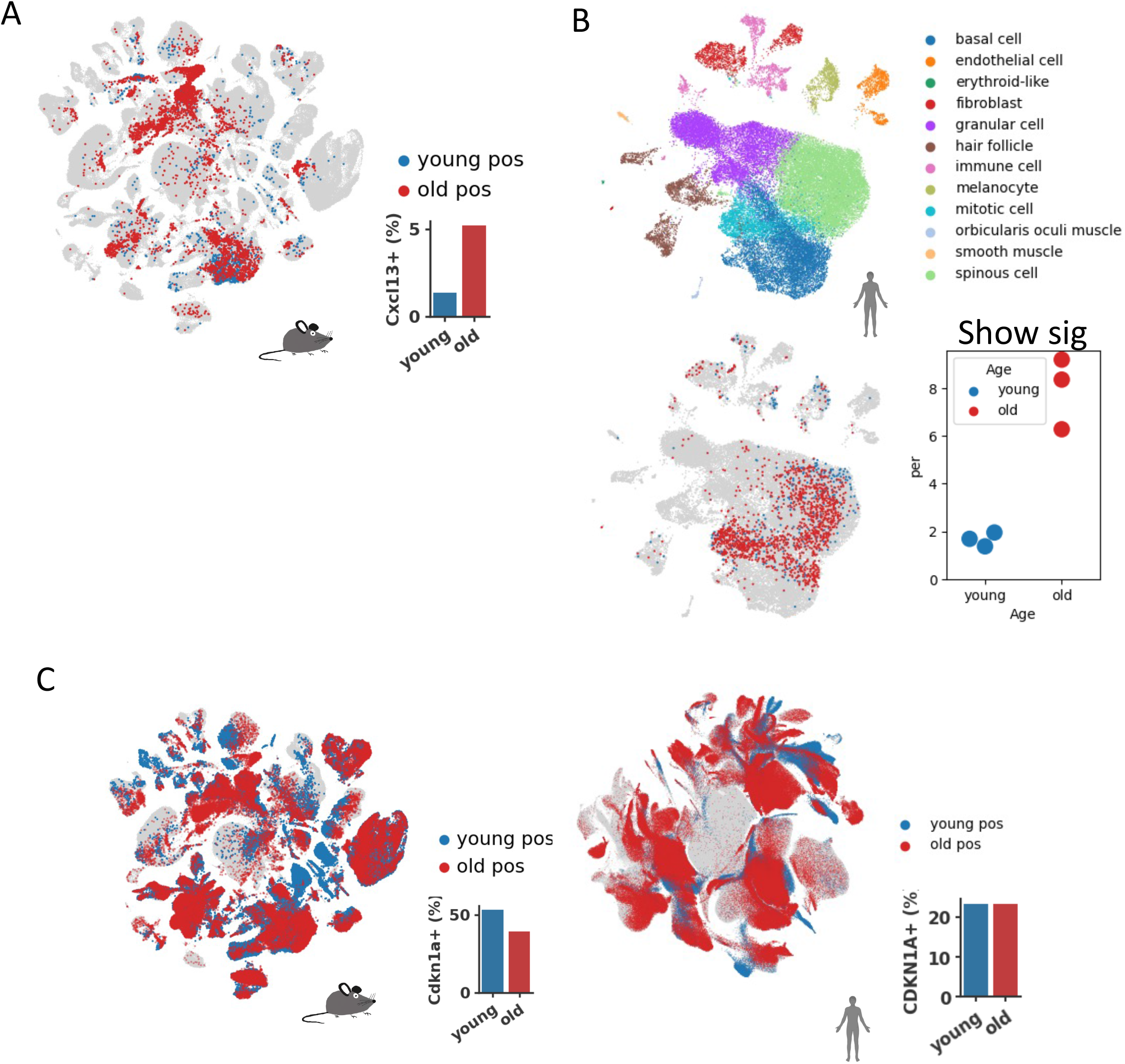
(**A**) UMAP representing *Cxcl13+* cells in all mouse cells. (**B**) UMAP (top) of human skin cells colored by cell type. UMAP (bottom) of human skill cells positive for *CDKN2A* from young (age 18-34) and old (age 34-58) donors. Dot plot depicts the percentage of *CDKN2A+* cells from old and young human skin cells. (**C**) UMAP representing *CDKN1A+* (gene that encodes p21^cip1^) cells in all mouse and human cells.

**Supplemental Figure 4.**
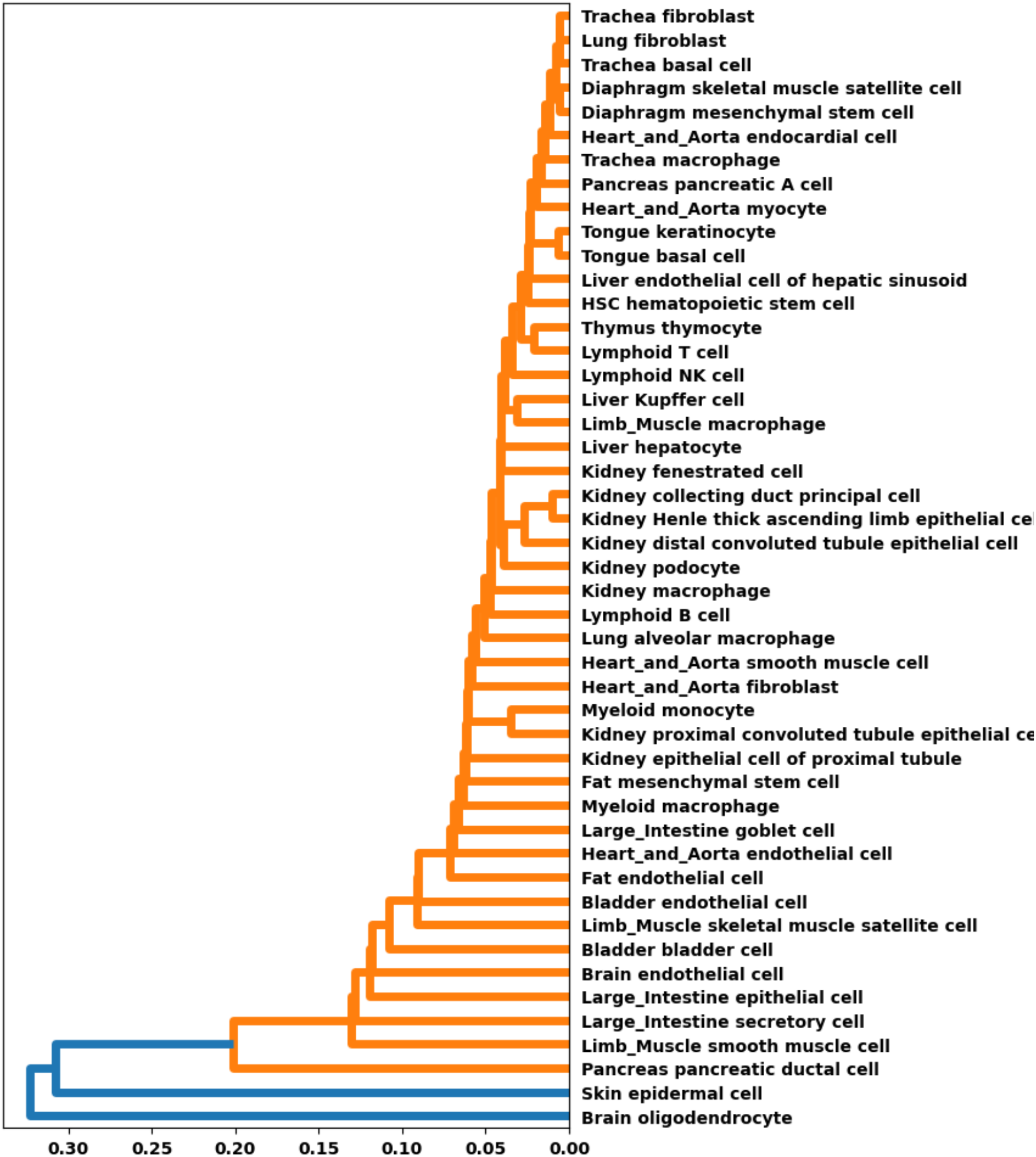
Hierarchal clustering of signatures based on gene-set similarity. Similarity is defined as the inverse of log_10_(BH-corrected Hypergeometric P) for each pairwise comparison

**Supplemental Figure 5.**
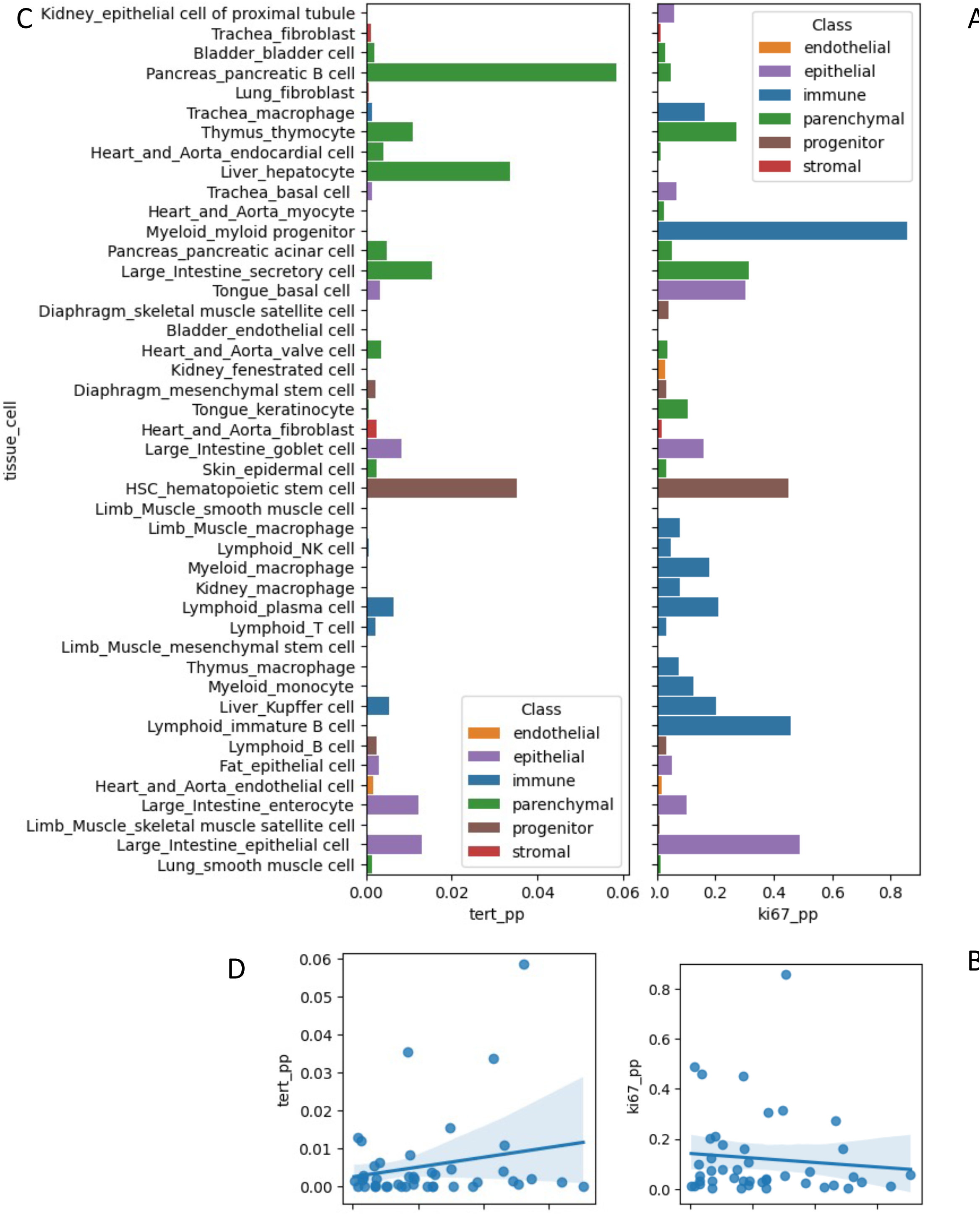
(**A**) The proportion of mouse cells expressing Tert and (**B**) Ki67. (**C,D**) The relationship between cell types expressing Tert and Ki67 with their gain in *SenePy* identified cells.

**Supplemental Figure 6.**
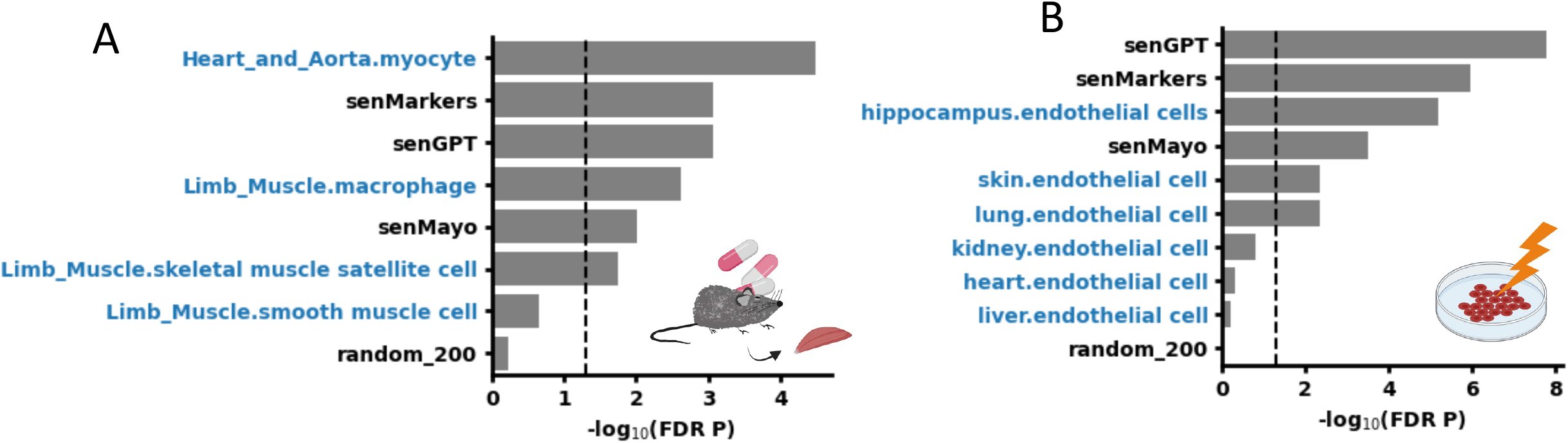
(**A**) Enrichment analysis of gene mRNAs less abundant in mouse muscle tissue after treatment with a senolytic. Blue lables indicate *SenePy* signatures. (**B**) Enrichment analysis of gene mRNAs more abundant after irradiation induced senescence of cultured endothelial cells.

